# Children exhibit greater persistence of motor learning-related patterns of hippocampal activity into post-task wake epochs

**DOI:** 10.64898/2026.04.02.716229

**Authors:** Anke Van Roy, Ainsley Temudo, Emily K. Taylor, Vincent Koppelmans, Kerstin Hödlmoser, Genevieve Albouy, Bradley R. King

## Abstract

Previous research has demonstrated that children exhibit superior - as compared to adults - consolidation of newly acquired motor sequences across post-learning periods of wakefulness. Given that consolidation is thought to be supported by the reactivation of learning-related patterns of brain activity during the rest periods following active task practice, we hypothesized that the childhood advantage in offline consolidation may be linked to greater reactivation during post-learning wakefulness. Twenty-two children (7-11 years) and 23 adults (18-30 years) completed two sessions of a motor sequence learning task, separated by a 5-hour wake interval. Multivoxel analyses of task-related and resting-state functional magnetic resonance imaging data were employed to assess the persistence of learning-related patterns of neural activity into post-task rest epochs, reflective of reactivation processes. Behavioral results demonstrated the previously reported childhood advantage in offline consolidation over a post-learning wake interval. Imaging results revealed that children exhibited greater persistence of task-related hippocampal – but not putaminal - activity into post-learning rest as compared to adults. These findings suggest that the childhood advantage in awake motor memory consolidation may be supported, at least partially, by enhanced reactivation of task-dependent hippocampal activity patterns during offline epochs.

## 1. Introduction

Behavioral evidence suggests that children exhibit superior offline (i.e., in the absence of active task practice) processing of recently learned movement sequences as compared to young adults (Adi-Japha et al., 2014; Ashtamker & Karni, 2013; Dorfberger et al., 2007; Van Roy et al., 2024; Van Roy & King, 2026). Specifically, whereas offline consolidation following a session of initial sequence learning is considered a slow, sleep-facilitated process in adults (see King et al., 2017 for review), pre-adolescent children appear to exhibit accelerated consolidation during post-learning wakefulness. Such accelerated consolidation is reflected by larger performance gains (Ashtamker & Karni, 2013; Van Roy et al., 2024) or greater resistance to interference from competing material (Adi-Japha et al., 2014; Dorfberger et al., 2007) emerging within minutes to hours following practice. This developmental advantage may not be limited to the timescale of hours following a learning session (i.e., “macro-offline” consolidation), as prior research has observed analogous results over “micro-offline” epochs (Du et al., 2017; Lagarrigue et al., 2025). More specifically, younger children (e.g., 6-year-olds) demonstrated greater improvements in performance during the short rest periods (i.e., on the timescale of minutes to seconds) commonly interspersed in between practice blocks as compared to older children (e.g., 10-year-olds) and adults (Du et al., 2017; Lagarrigue et al., 2025; but see Van Roy et al., 2024).

Investigations into the neural correlates of motor sequence learning in children, and the developmental advantage in offline processing in particular, are scarce [but see Hedenius & Persson, 2022; Thomas et al., 2004]. We recently conducted a neuroimaging study that aimed to address this knowledge gap (Van Roy et al., 2025). Behavioral results replicated the childhood advantage in awake macro-offline consolidation, whereas there was no clear evidence for differences in micro-offline performance gains between children and adults. Neuroimaging results revealed that children exhibited smaller modulations in brain activity between task and rest epochs in a widespread network of brain regions, including the sensorimotor cortex, supplementary motor area, cerebellum, putamen and regions commonly associated with the default mode network. Moreover, similar levels of activity during task and rest epochs in the hippocampus as well as the dorsolateral prefrontal and somatosensory cortices were associated with better macro-offline consolidation in children. Interestingly, recent research in adults demonstrated that higher levels of activity during rest (relative to task practice) were related to the persistence of task-related activity patterns into subsequent rest epochs, with this persistence reflecting the reactivation of learning-related neural activity (Gann et al., 2023). Our findings in children thus potentially suggest that the continued engagement of the developing brain during offline rest may reflect the reactivation of learning-related brain activity.

Post-learning reactivation or replay was originally demonstrated in seminal animal work (Griffin et al., 2025; Pavlides & Winson, 1989; Wilson & McNaughton, 1994) and an analogous pattern of results was later observed in humans after declarative and perceptual learning (Hermans et al., 2016; Schuck & Niv, 2019; Tambini & Davachi, 2013, 2019; Wittkuhn & Schuck, 2021). Similarly, research from our team and others in the motor memory domain demonstrated that both micro- and macro-offline consolidation processes are supported by neural reactivation (Buch et al., 2021; Gann et al., 2023; King et al., 2022). Specifically, this research suggested that patterns of brain activity observed during active task practice (i.e., online learning) are spontaneously reactivated or replayed during subsequent offline epochs, leading to a strengthening of the recently acquired memory trace (Buch et al., 2021). This task-related reactivation was demonstrated in the adult hippocampus and putamen (Buch et al., 2021; Gann et al., 2023; King et al., 2022), two brain regions known to be critical for motor learning and memory consolidation processes in healthy adults (see Albouy et al., 2013 and King et al., 2017 for reviews). Based on this prior work, we hypothesized that the childhood advantages in offline motor memory processing at different timescales would be attributed to greater reactivation of task-related activity patterns in the hippocampus and putamen during inter-block and post-learning rest epochs. To test this possibility, the current study employed a multivoxel analysis to characterize the similarity of brain activity patterns between task and both micro- and macro-offline rest epochs, an analytic approach thought to reflect reactivation processes.

## 2. Methods and materials

Data collection procedures are outlined in the first publication from the same overarching research project (Van Roy et al., 2025), where complete details on the full protocol can be found. Here, we provide methodological information pertinent to the objective outlined above. Data collection and analysis plans were registered in April 2024 via Open Science Framework (https://doi.org/10.17605/OSF.IO/SW6D8). Any additional collection or analytic procedures that were not included in the registration are labeled as exploratory in this manuscript. Any deviations from the registered protocol are marked in this text with an asterisk and explained in Appendix 1. The behavioral and imaging data as well as the analytic pipelines will be made publicly available on Zenodo upon journal publication. The study protocol was approved by the University of Utah Ethics Committee (IRB_00155118).

### 2.1 Participants

Six- to 12-year-old children and 18-30-year-old adults of all genders were recruited via advertisements posted in public spaces and on relevant websites. Adult participants and parents of child participants gave informed consent, and children between 7 and 12 years of age also provided informed assent. All participants were right-handed, reported no known history of medical, neurological, psychological, or psychiatric conditions (including depression and anxiety), were free of psychotropic or psychoactive medications, indicated no abnormal or irregular sleep, showed no MRI contraindications (including claustrophobia), and had no mobility limitations preventing successful motor task completion (all criteria assessed based on self- or parental-report). Furthermore, none of the participants received extensive training in a musical instrument requiring dexterous finger movements (e.g., piano) or as a typist, nor had they previously participated in a research experiment employing a similar motor sequence learning task. According to the sample size estimation outlined in our registration and in Van Roy et al. (2025), the required sample size when assuming a power (1-beta) of 0.8 and an alpha level of 0.05 to detect a childhood advantage in awake motor memory consolidation consisted of 46 participants (23 children; 23 adults). Sixty-one participants initiated participation; the final sample consisted of 22 children (14 females, 7 males and 1 non-binary; mean age = 9.5 years) and 23 adults (16 females, 6 males and 1 non-binary; mean age = 23.4 years; see Van Roy et al., 2025 for reasons for exclusion of participants). Note that although 6- to 12-year-old children were recruited, the final sample was limited to children between 7 and 11 years of age. Moreover, small subsets of participants were excluded from specific statistical contrasts presented in this manuscript. The reasons for any exclusions from specific analyses are outlined in Appendix 1 and the sample sizes included in analysis are explicitly stated in the corresponding figure or table captions.

### 2.2 Experimental procedure

The full study protocol consisted of an initial familiarization visit followed by two experimental sessions that took place on the same day (see Figure 1). For the nights prior to both testing days, participants were instructed to get a good night of sleep (i.e., bedtime no later than 1 am and at least 7 hours of sleep). The familiarization session (∼1 hour) focused on habituating the participants to the research team, MR environment using a mock scanner and our series of motor tasks. A detailed description of the familiarization procedure is provided in the supplemental material of our previous publication (Van Roy et al., 2025).

**Figure 1.**
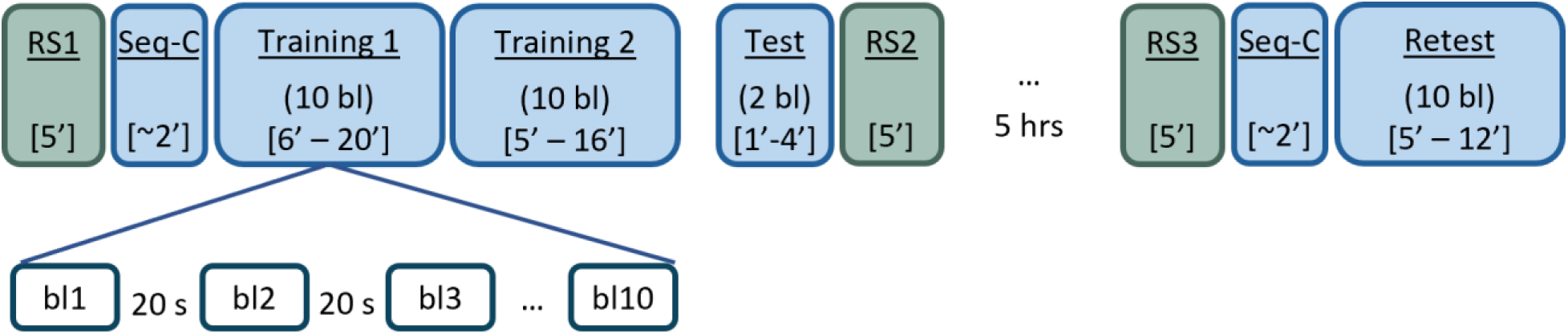
Experimental design. Initial acquisition of the motor task consisted of training 1, training 2 and a post-learning test. Resting-state (RS) scans were acquired prior to and following the FTT task runs (RS1 and RS2, respectively). Following a 5-hour macro-offline consolidation interval, participants completed another RS scan (RS3), followed by a retest of the motor sequence. hrs = hours. Seq-C = sequence-check. bl = block. Durations of runs are indicated between brackets (units of minutes). Note that ranges were provided for task runs since durations depended on the individual’s performance. BOLD images were acquired during RS and all FTT task runs, except for the sequence-checks. Note that the familiarization visit was not included in the figure, as these data were not analyzed.

The primary experimental day consisted of two sessions that were completed approximately 5 hours apart. The first session, referred to as training, took place between 9 am – 1:30 pm. The second session was a 5-hour delayed retest that took place between 2 pm – 6:30 pm and afforded the assessment of motor memory consolidation following an offline interval of wakefulness. During the initial training session, participants first completed questionnaires, an objective assessment of vigilance (i.e., the Psychomotor Vigilance Task - PVT; Dinges & Powell, 1985; see Van Roy et al., 2025 for results) and further familiarization with the motor tasks. Subsequently, participants were positioned in the 3-Tesla MR scanner with an MR compatible keyboard on each hand. Following a pre-learning resting-state run (RS1), participants completed a short sequence-check run in which they performed the explicitly-provided sequence (4-2-1-3-4-1, where 1 is the left middle finger, 2 is the left index finger, 3 is the right index finger and 4 is the right middle finger) slowly and as accurately as possible until three correct sequences were performed consecutively. Participants then completed initial learning of the motor sequence (finger tapping task - FTT; see section 2.3), consisting of two training runs and a test phase, while blood-oxygen-level-dependent (BOLD) images were acquired. The training portion was decomposed into two 10-block runs to provide participants with a rest opportunity during which they could interact with the research team. The test run consisted of 2 blocks and was administered approximately 1 minute after the training portion, affording the assessment of end-of-training performance following the dissipation of fatigue (Pan & Rickard, 2015). BOLD images were acquired during the post-learning test to mimic other runs; however, the corresponding MRI data were not analyzed. A post-learning resting-state run (RS2) concluded the initial learning session. Participants then left the imaging facility and were instructed to avoid strenuous exercise, caffeine, sleep, and practicing the motor task.

Approximately 5 hours after the end of the training session, participants returned for the delayed retest. Following brief questionnaires and the PVT, participants were positioned back in the MR scanner. The retest protocol consisted of a pre-retest RS run (RS3) and the sequence-check, followed by 10 blocks of the FTT. During all 3 RS scans, participants were instructed to focus on a white cross on a dark screen and to not fall asleep nor think of anything in particular for the duration of the scan.

### 2.3 Finger Tapping Task (FTT)

A version of the explicit, bimanual finger tapping task (FTT) - used previously by our team (e.g., Dolfen et al., 2019) - was modified for children and used to assess motor sequence learning and memory consolidation. In brief, participants used the index and middle fingers of both hands to perform a 6-element sequence of keypresses (i.e., 4-2-1-3-4-1). Participants were asked to repeatedly tap the finger sequence on the keyboard as rapidly and accurately as possible while a green cross and the sequence of numbers were displayed on the screen. Each practice block consisted of 48 keypresses (i.e., ideally corresponding to 8 repetitions of the 6-element sequence) and ended once this number of keypresses was recorded. Practice blocks alternated with 20-second rest blocks, during which the cross turned red and the sequence of numbers was replaced by a fixed sequence of random symbols (i.e., @-!-$-%-&-#) to avoid exposure to the sequence during rest. Participants were instructed to focus on the red cross and to not move their fingers during these rest epochs. The timing and number of each keypress were recorded during performance of the FTT.

### 2.4 MRI data acquisition

MRI data were acquired on a Siemens MAGNETOM Vida 3.0T MRI system equipped with a 64-channel head coil. To minimize head motion, additional padding was placed around the head. For each participant, a high-resolution, T1-weighted (T1w) anatomical image was acquired with a 3D MPRAGE sequence (axial; repetition time (TR) = 2500 ms; echo time (TE) = 2.98 ms; inversion time (TI) = 1070 ms; flip angle (FA) = 8°; 176 slices; field-of-view (FoV) = 256 x 256 x 176mm3; voxel size = 1.0 x 1.0 x 1.0 mm3; duration = 5:59 min). Parallel acquisition was conducted in the GRAPPA mode, with reference line phase encoding (PE = 32), and acceleration factors PE and 3D equal to 2 and 1, respectively.

Resting-state (RS) and task-related fMRI data were acquired with an interleaved gradient EPI pulse sequence for T2*weighted images (SMS acceleration with multiband factor 5; TR = 797.0 ms; TE = 31.0 ms; flip angle = 59°; 55 transverse slices; voxel size = 2.5 x 2.5 x 2.5 mm3; field of view = 220 x 220 x 138 mm3). The RS runs consisted of 375 acquired volumes with a duration of 5:06 minutes (including unsaved dummy pulses). The task-related runs were set to a max of 1500 volumes, with a maximum duration of 20:03 minutes (including unsaved dummy pulses). The exact number of scans in each of the task runs depended on the individual’s FTT performance speed.

Gradient-recalled echo (GRE) fieldmaps were acquired to correct for distortions due to magnetic field inhomogeneities. Specifically, two phase-encoded GRE images with different echo times (TR = 752 ms; TE1 = 5.19 ms, TE2 = 7.65 ms; flip angle = 90°; 55 transverse slices; voxel size = 2.5 × 2.5 × 2.5 mm3; field of view = 220 × 220 × 138 mm3) were collected prior to task performance during the training and retest sessions. Phase-difference maps were calculated from the two GRE images using in-scanner processing.

### 2.4 Data processing

The sections below detail the preprocessing of behavioral and neural data, as well as statistical testing. All statistical tests were performed using R version 4.5.1 (Posit, PBC, Boston, Massachusetts, USA). For behavioral data, alpha was set to 0.05. For the sake of completeness, those effects when 0.05 ≤ p ≤ 0.1 are also highlighted as non-significant trends. For the multivoxel correlation structure (MVCS) analyses of the imaging data, results in the main text were Bonferroni corrected for the two primary regions of interest (ROIs; see below for justifications for these specific regions), changing the alpha level for significance to 0.025. In the event of a sphericity violation, a Greenhouse-Geisser correction was applied. Depending on the statistical test, eta-squared, partial eta-squared or Hedge’s G were reported as effect size measures. In addition to null hypothesis testing, Bayes Factors (BF; anovaBF, ttestBF and regressionBF functions from the BayesFactor package version 0.9.12-4.7 in R) were computed to assess the likelihood of the observed data favoring an alternative model (e.g., evidence for differences among group, blocks, etc.) relative to the null model (i.e., no differences).

#### 2.4.1 Behavioral analyses

A comprehensive characterization of the behavioral results was provided in Van Roy et al. (2025). Here, we simply focus on the details of our behavioral metrics that will be related to the persistence of learning-related brain activity patterns into subsequent rest intervals (i.e., offline changes in performance). For each FTT practice block, performance speed was computed as the median time to perform a transition between pairs of consecutive keypresses. Performance accuracy was calculated as the percentage of consecutive keys that were pressed in the same order as presented in the sequence (i.e., correct transitions). Consistent with our registration and to account for expected age group differences independent of learning (Meulemans & Van der Linden, 1998; Salehi et al., 2016; Sugawara et al., 2014), outcome measures were normalized relative to the outcome measures of the first training block.

For the assessment of macro-offline consolidation, performance changes (both speed and accuracy metrics) across the 5-hour post-learning period of wakefulness were computed as the difference in normalized performance from the end of training (average across the 2 post-learning test blocks) to the beginning of retest (average across the first 2 blocks). As outlined in our registration, age group differences in macro-offline performance changes were assessed using one-tailed independent samples t-tests. Micro-offline performance changes were computed as the difference between the median normalized transition times of the last 6 transitions of one practice block and the first 6 transitions of the subsequent block (i.e., across the short rest periods; last 6 transitions_n_ – first 6 transitions_n+1_, with n = practice block). Micro-offline performance changes were averaged across rest intervals to obtain a single value per training run and were compared between age groups and training runs using a group (i.e., children versus adults) by training run (i.e., training 1 versus training 2) ANOVA.

#### 2.4.2 MRI preprocessing

The DICOM images output by the SIEMENS scanner were converted to BIDS using the dcm2bids tool, version 3.1.1 (Boré et al., 2023), implemented in Python 3.11.5. Prior to any preprocessing, the coordinate system in the headers of the T1-weighted images were reoriented to the Montreal Neurological Institute (MNI) template with images remaining in native space.

Initial spatial preprocessing was performed in fMRIPrep 24.1.0* (Esteban et al., 2019). Appendix 2 contains a description of the spatial preprocessing that was automatically generated by fMRIPrep and released under the CC0 license. Here, we provide a brief overview of key preprocessing steps. The T1w anatomical image was corrected for intensity non-uniformity, skull-stripped and segmented into white matter (WM), gray matter (GM) and cerebrospinal fluid (CSF). For each task-related and resting-state run, head motion was estimated (see Appendix 3 for information on the magnitude of head motion parameters and corresponding statistical comparisons), and functional images were corrected for susceptibility distortion using GRE fieldmaps as well as co-registered to anatomical space. Several confounding time-series were calculated based on the preprocessed BOLD data, such as framewise displacement (FD), and anatomical noise components derived from CSF and WM probabilistic maps were identified using principal component analysis (PCA; explaining >50 % of variance; see below for details on preprocessing steps that used these confounding time-series). Consistent with previous research employing similar multivoxel analytic approaches (King et al., 2022; Tambini & Davachi, 2013), anatomical and functional images remained in native space (i.e., not normalized to a template).

Additional preprocessing of the MRI images was performed in Matlab R2023a, utilizing SPM12 (version r7771) functions (Welcome Department of Imaging Neuroscience, London, UK). The BOLD signal from specific regions of interest (ROIs) was extracted for all epochs of interest (resting-state runs, blocks of active task practice and interleaved rest periods; see Appendix 3 for the number of voxels included per ROI). The time series extracted from the voxels of each ROI was then subjected to detrending, high-pass filtering (cutoff = 1/128), and the scrubbing of volumes with excessive head motion (FD > 0.5 frames, including 1 frame forward and 1 backward; using the head motion measures output by fMRIPrep). Furthermore, volumes with maximum translation > 5mm were excluded along with all subsequent volumes if the excessive movement occurred toward the end of the run, or all preceding volumes if the head movement happened toward the beginning (see Appendix 1 for details). Lastly, a subset of confounding time series computed in fMRIPrep was used for nuisance regression. These variables included head motion estimates (i.e., translation and rotation across all three axes) and their temporal derivatives and quadratic terms, as well as the top 5 components of the PCA conducted on signals from the CSF and WM masks.

#### 2.4.3 Multivoxel Correlational Structure (MVCS) analysis

Consistent with our registration, the primary regions of interest (ROIs) for the MVCS analyses were the bilateral hippocampus and putamen. The bilateral nucleus accumbens served as the control region of no interest (i.e., thought to *not* play a role in motor memory). Results from these 3 registered ROIs (two primary and one control) are reported in the main text. For completeness, exploratory (i.e., not registered) analyses were performed for six cortical (i.e., primary motor cortex, pre-supplementary motor area, supplementary motor area, ventral premotor area, dorsal premotor area, primary sensory cortex) and one subcortical region of interest (i.e., caudate nucleus). These exploratory regions were included based on previous research linking these regions to motor sequence learning and memory consolidation processes [e.g., Berlot et al., 2020; Gann et al., 2023b; Temudo et al., 2025]. Masks for all subcortical regions were created using the FMRIB’s Integrated Registration Segmentation Toolkit (FIRST; FSL First, Oxford University, Oxford UK; Patenaude et al., 2011), while masks for cortical regions were extracted using the Human Motor Area Template (HMAT; Mayka et al., 2006) and subsequently transformed into native space using the individual’s inverse deformation field output from segmentation of the anatomical image. An additional exclusion mask was applied to these ROIs to eliminate voxels located outside the brain or with a grey-matter probability below 10%.

The multivoxel correlational structure analysis was similar to our previous research (Gann et al., 2021, 2023; King et al., 2022). Following the preprocessing described above, correlations among the n-voxels extracted from each ROI were computed for each epoch of interest (see Figure 2). Specifically, we calculated the Pearson correlation between each of the n BOLD-fMRI time courses, yielding an n-by-n MVCS matrix that reflects the pattern of brain activity in the ROI during that specific epoch. The Pearson correlation coefficients were Fisher Z-transformed to ensure normality. Epochs of interest included (a) pre-learning resting-state (i.e., RS1), (b-c) blocks of active task practice during training runs 1 and 2, (d-e) inter-block rest periods in training runs 1 and 2, and (f) post-learning resting-state (i.e., RS2). For the blocks of active task practice and periods of inter-block rest, the BOLD signal was extracted from those specific epochs and concatenated into one continuous signal. Next, similarity indices (SIs) were computed, reflecting the similarity of the patterns of brain activity between pairs of epochs of interest. Specifically, the r-to-z transformed correlations between the below-diagonal elements of the MVCS matrices from two epochs of interest were computed (King et al., 2022; Tambini & Davachi, 2013). The specific epochs used in the computation of SIs depend on the research question being addressed and are outlined in Section 2.4.4 below.

**Figure 2.**
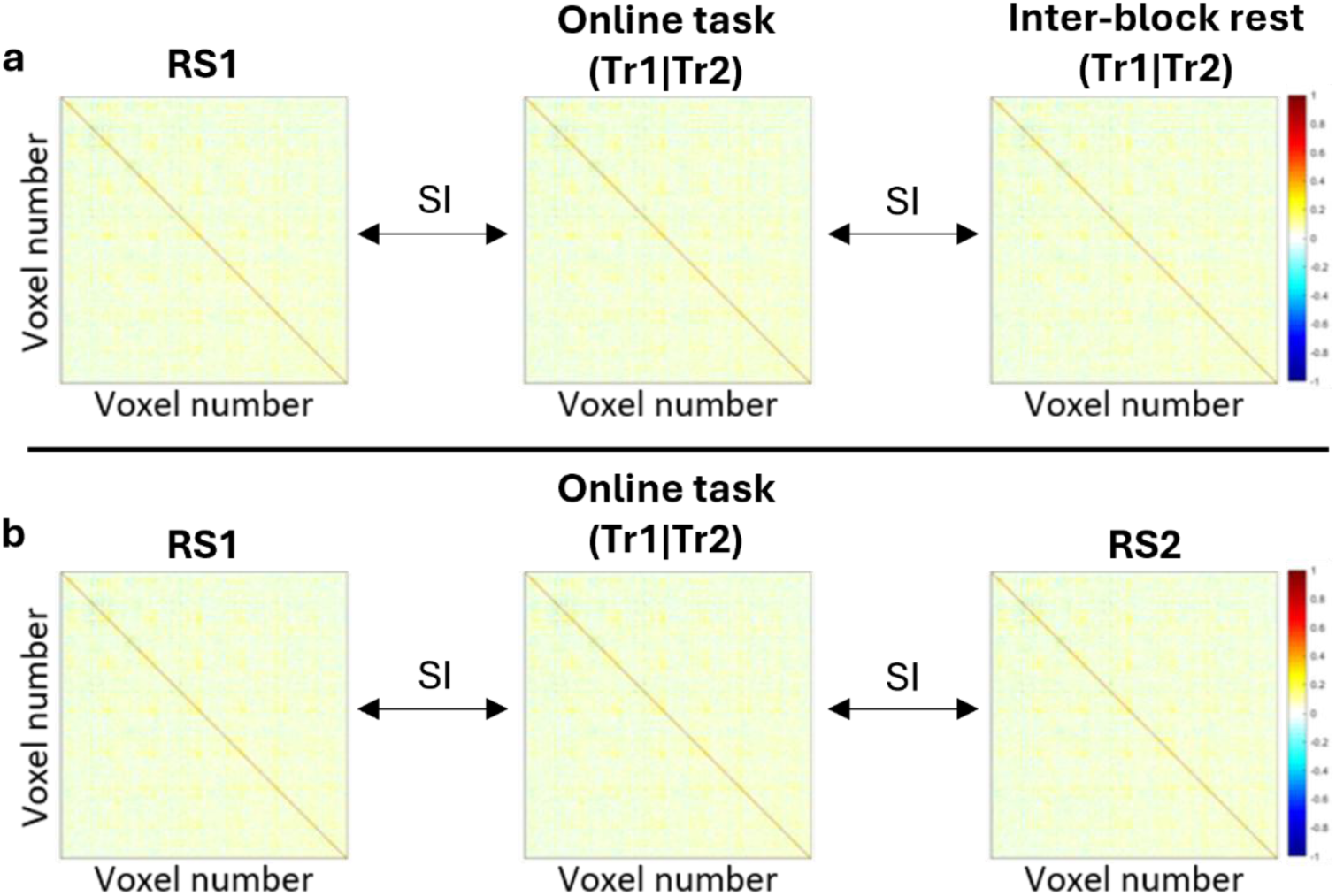
MVCS matrices for an exemplar ROI and participant for the assessment of **(a)** inter-block and **(b)** post-learning persistence. Similarity indices (SI) between online task practice of each training run and the rest epoch of interest (i.e., either inter-block intervals or the post-learning RS2 scan) were computed, as well as between online task practice and the pre-learning RS1 scan. Normalized SIs were obtained by subtracting the SIs between online task practice and RS1 from the SIs between online task practice and the rest epoch of interest (see Section 2.4.4 for details).

#### 2.4.4 Statistical analyses of MVCS data

Prior to assessing task-related persistence into offline epochs, and consistent with our registration, we first assessed whether the age groups differed in the similarity between activity patterns during online task practice of each training run and pre-learning rest (i.e., RS1). As significant differences between children and adults were observed (see Appendix 2), age group differences in the similarity in activity patterns between online task practice and baseline rest were accounted for in the further analyses (see below) of all studied regions*.

Age group differences in the persistence of task-related patterns of activity into inter-block rest, thought to reflect micro-offline reactivation processes, were assessed by computing SIs between the online task practice and inter-block rest MVCS matrices within each training run. As a within-subject control and similar to previous research (King et al., 2022; Tambini & Davachi, 2013), SIs were also computed between online task practice (both runs) and the pre-learning RS run (RS1). Next, a normalized inter-block persistence measure was obtained for each training run by subtracting the SI between online task practice and RS1 from the SI between online task practice and inter-block rest of the same training run (i.e., SI (online practice | inter-block rest) – SI (RS1 | online practice)). Positive values reflect a greater similarity in activity patterns between online task practice and inter-block rest as compared to pre-learning rest. The normalized inter-block persistence measure was then subjected to a group (children versus adults) by training run (training 1 versus training 2) ANOVA. Finally, the relationship between inter-block persistence and behavioral micro-offline performance changes was assessed using separate multiple linear regressions with the normalized inter-block persistence measure as the independent variable and the micro-offline performance changes as the dependent variable. An age group (dummy coded) by normalized SI interaction variable was included in the regression model to examine whether the relationship differed between the age groups.

Next, we examined age group differences in the persistence of task-related activity patterns into post-learning rest (i.e., reflecting reactivation on the macro-offline timescale). Specifically, we computed the SIs between online task practice (both runs) and the post-training RS run (RS2), as well as between online task practice of each run and RS1 (i.e., baseline rest; used as within-subject control). Similar to above, a normalized post-learning persistence measure was obtained for each training run by subtracting the SI between RS1 and the respective online task practice from the SI between RS2 and the same online task practice (i.e., SI (online practice | RS2) – SI (RS1 | online practice)). Larger values (i.e., more positive) are indicative of greater persistence of the task-related activity patterns into post-learning rest. The normalized post-learning persistence measure was then entered into a mixed ANOVA with group (2 levels: children versus adults) as a between-subject factor and training run (2 levels: training 1 versus training 2) as within-subject factor. Lastly, the relationship between post-learning persistence and behavioral macro-offline performance changes was examined using multiple linear regression analyses. Specifically, separate multiple linear regressions were conducted for each training run with the normalized persistence measure as the primary independent variable and the behavioral macro-offline performance changes as the dependent variable. An age group (dummy coded) by normalized persistence interaction variable was included in the regression model.

As a follow-up to the group differences in post-learning persistence (see results in section 3.2.2), we conducted an exploratory analysis that examined whether persistence of task-related activity into RS2 was still present during the post-consolidation resting-state run (i.e., RS3). Specifically, SIs were computed between online task practice of training 2 and the post-consolidation RS run (RS3), as well as between online task practice of training 2 and RS1 (i.e., baseline rest; used as within-subject control). Similar to above, a normalized post-consolidation persistence measure was obtained by subtracting the SI between RS1 and online task practice from the SI between RS3 and online task practice (i.e., SI (online practice | RS3) – SI (RS1 | online practice)). Larger values (i.e., more positive) are indicative of greater persistence of the task-related activity patterns into post-consolidation rest (i.e., RS3). The normalized post-consolidation persistence measure was then entered into a group (i.e., children vs. adults) by RS run (i.e., RS2 vs. RS3) ANOVA.

The inter-block and post-learning persistence analyses outlined above compare differences in offline persistence between the two age groups. Yet, to provide more fine-grained analyses of developmental changes, we performed exploratory analyses assessing age-related changes in persistence across childhood. Specifically, the normalized inter-block and post-learning persistence measures were subjected to separate regression analyses with age as a continuous independent variable. For both dependent measures, five potential fit options (i.e., single exponential, double exponential, linear, quadratic and power functions) were tested. The final model was selected based on standard assessments of model fit (i.e., AIC). Results regarding these age-related changes during childhood are provided in Appendix 4.

## 3 Results

### 3.1 Behavioral results

A detailed characterization of the behavioral results, as well as participant characteristics, sleep leading up the experimental sessions and vigilance at the time of testing, are provided in Van Roy et al. (2025). Here, we focus on the behavioral metrics reflecting offline consolidation processes that may be linked to persistence of task-related brain activity patterns.

Results revealed a non-significant trend toward larger micro-offline performance changes in children, although the Bayes Factor suggested weak evidence in support of no group differences (F_(1,43)_ = 3.70, p = 0.061, ƞ^2^ = 0.044, BF_10_ = 0.869; see Figure 3, panel a). Training run 1 exhibited larger micro-offline performance changes as compared to training run 2 (F_(1,43)_ = 7.99, p = 0.007, ƞ^2^ = 0.080, BF_10_ = 12.154), a difference that was comparable between age groups (F_(1,43)_ = 0.00, p = 0.963, ƞ^2^ < 0.001, BF_10_ = 0.287).

**Figure 3.**
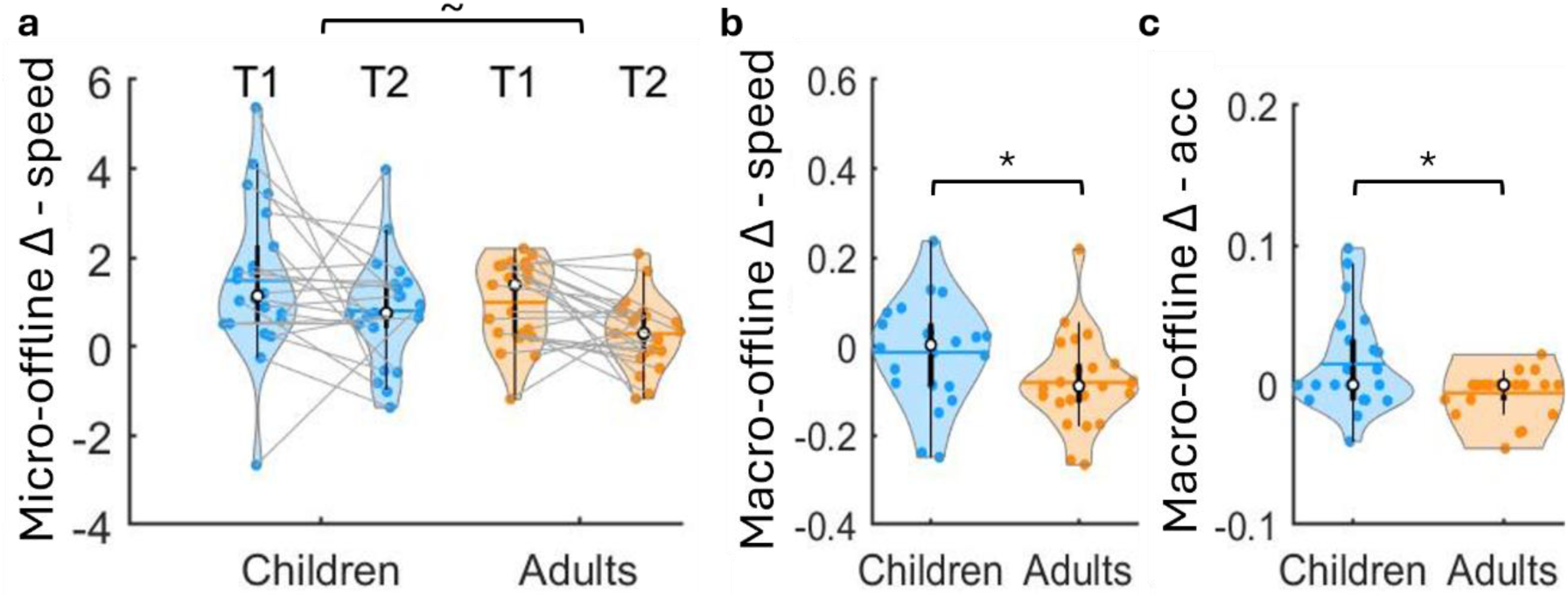
Micro- and macro-offline performance changes. **(a)** Micro-offline performance changes summed across blocks for training runs 1 and 2 (T1 and T2, respectively). n = 22 children for T1 and 21 for T2. n = 23 adults. **(b-c)** Macro-offline changes in normalized speed (**b**) and accuracy (**c**) across the 5-hour offline period for children and adults. n = 22 children and 23 adults. Positive values are indicative of performance improvements. Shaded regions represent the kernel density estimates of the data, colored circles depict individual data, open circles represent group medians, and the horizontal lines depict group means (Bechtold et al., 2021). * p < 0.05 and ∼ p < 0.1 for group differences.

The assessment of the consolidation process across the macro-offline interval (see Figure 3, panels b and c) found significantly greater macro-offline changes in children than adults for both speed (t_43_ = 2.059, p = 0.023, G = 0.60, BF_10_ = 1.573) and accuracy (t_28.685_ = 2.527, p = 0.009, G = 0.74, BF_10_ = 3.812). As reported in our original paper with these data (Van Roy et al., 2025), these results confirm the behavioral childhood advantage in offline processing previously shown by our team (Van Roy et al., 2024) and prior studies (Adi-Japha et al., 2014; Ashtamker & Karni, 2013; Dorfberger et al., 2007).

### 3.2 MVCS results

#### 3.2.1 Persistence of task-related activity into inter-block rest epochs

Normalized inter-block persistence, reflecting reactivation on the micro-offline timescale, for each age group and training run is depicted in Figure 4. Results revealed a significant effect of group for the hippocampus (HC; F_(1,38)_ = 18.04, p < 0.001, ƞ^2^ = 0.281, BF_10_ = 332.143) and putamen (PUT; F_(1,38)_ = 8.31, p = 0.006, ƞ^2^ = 0.146, BF_10_ = 6.774), but not the nucleus accumbens control region (ACCU; F_(1,38)_ = 0.13, p = 0.719, ƞ^2^ = 0.003, BF_10_ = 0.403). More specifically, children showed greater inter-block persistence of task-related patterns of activity in the two task-relevant regions as compared to adults. There were no significant differences between training runs for any studied region, as evidenced by the absence of a training run effect (HC: F_(1,38)_ = 0.41, p = 0.526, ƞ^2^ = 0.002, BF_10_ = 0.261; PUT: F_(1,38)_ = 0.04, p = 0.837, ƞ^2^ < 0.001, BF_10_ = 0.227; ACCU: F_(1,38)_ = 1.12, p = 0.297, ƞ^2^ = 0.006, BF_10_ = 0.420) and group x training run interaction (HC: F_(1,38)_ = 0.02, p = 0.887, ƞ^2^ < 0.001, BF_10_ = 0.285; PUT: F_(1,38)_ = 0.05, p = 0.829, ƞ^2^ < 0.001, BF_10_ = 0.377; ACCU: F_(1,38)_ = 0.03, p = 0.873, ƞ^2^ < 0.001, BF_10_ = 0.310). It is worth noting that similar results were found in the exploratory regions of interest (see Appendix 4), including the caudate nucleus, primary motor cortex (M1), pre-supplementary motor area (pre-SMA), supplementary motor area (SMA), ventral and dorsal premotor cortices (PMv and PMd) and primary somatosensory cortex (S1). Altogether, children exhibited greater inter-block persistence of task activity patterns in all task-relevant brain regions, including our primary ROIs the hippocampus and putamen, but not in the nucleus accumbens control region.

**Figure 4.**
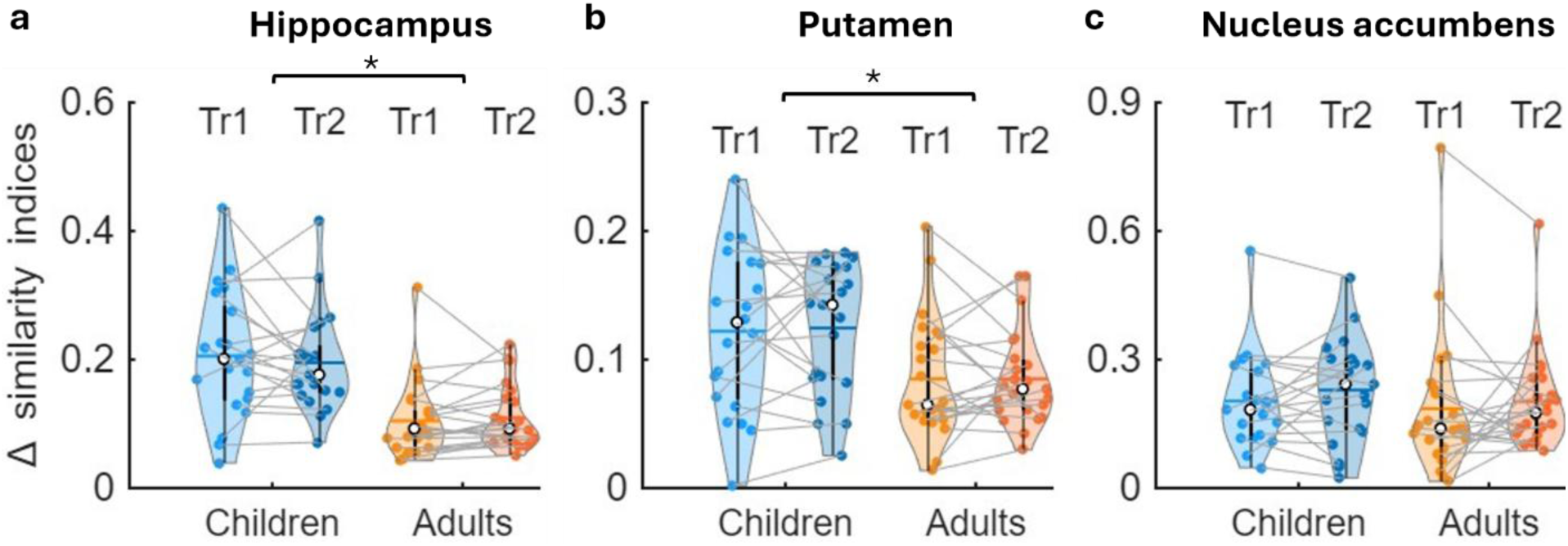
Normalized inter-block persistence for the **(a)** hippocampus, (**b**) putamen, and (**c**) nucleus accumbens. Shaded regions represent the kernel density estimates of the data, colored circles depict individual data, open circles represent group medians, and the horizontal lines depict group means (Bechtold et al., 2021). * p < 0.05 for group effects. Tr1 = training 1; n = 18 children, 23 adults. Tr2 = training 2; n = 19 children, 23 adults. Please note that the y-axes scale changes across the 3 panels.

#### 3.2.2 Persistence of task-related activity into post-learning rest

Normalized post-learning persistence, reflecting reactivation during the early macro-offline consolidation interval, for each age group and training run are displayed in Figure 5. Results revealed no significant group x training run interaction for any of the brain regions (HC: F_(1,41)_ = 0.05, p = 0.818, ƞ^2^ < 0.001, BF_10_ = 0.314; PUT: F_(1,41)_ = 0.06, p = 0.809, ƞ^2^ < 0.001, BF_10_ = 0.306; ACCU: F_(1,41)_ = 0.96, p = 0.332, ƞ^2^ = 0.009, BF_10_ = 0.417). There was a significant difference between training runs for all studied regions, with patterns of activity during the post-learning resting-state scan being significantly more similar to Training 2 as compared to Training 1 (HC: F_(1,41)_ = 15.21, p < 0.001, ƞ^2^ = 0.136, BF_10_ = 251.002; PUT: F_(1,41)_ = 33.79, p < 0.001, ƞ^2^ = 0.125, BF_10_ = 26081.03; ACCU: F_(1,41)_ = 17.57, p < 0.001, ƞ^2^ = 0.142, BF_10_ = 663.003). Additionally, children showed significantly greater post-learning persistence, across training runs, as compared to adults for the hippocampus (F_(1,41)_ = 6.92, p = 0.012, ƞ^2^ = 0.089, BF_10_ = 1.802), but not the putamen (F_(1,41)_ = 0.38, p = 0.542, ƞ^2^ = 0.008, BF_10_ = 0.362), nucleus accumbens (F_(1,41)_ = 0.76, p = 0.387, ƞ^2^ = 0.011, BF_10_ = 0.333) or any of the exploratory regions (see Appendix 4).

**Figure 5.**
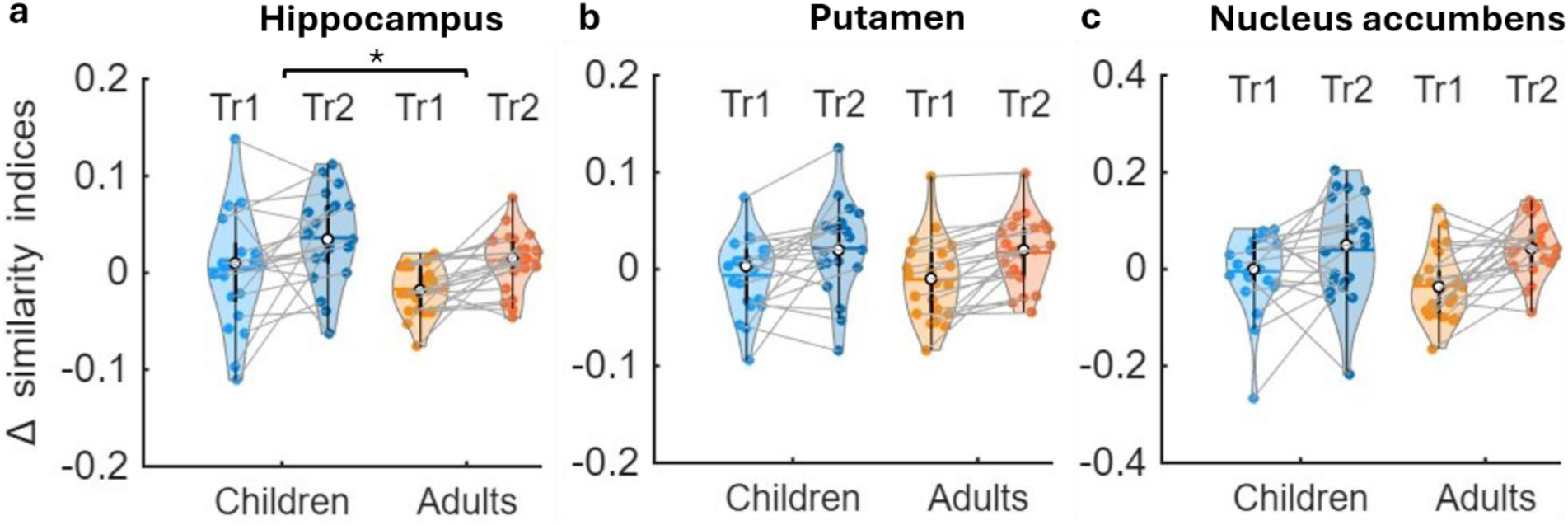
Post-learning persistence depicted for the (**a**) hippocampus, (**b**) putamen, and (**c**) nucleus accumbens. Shaded regions represent the kernel density estimates of the data, colored circles depict individual data, open circles represent group medians, and the horizontal lines depict group means (Bechtold et al., 2021). * p < 0.05 for group effects. n = 20 children and 23 adults. Please note that the y-axes scale changes across the 3 panels.

As a follow-up to the age group differences in hippocampal post-learning persistence, we conducted an exploratory analysis that examined whether the greater persistence of task-related hippocampal activity into the RS2 observed in children was still present following the 5-hour offline consolidation interval (i.e., observed in RS3; see Appendix 4). Results revealed a significant group x RS run (i.e., RS2 vs. RS3) interaction effect (F_(2,83.85)_ = 4.39, p = 0.015, ƞ^2^ = 0.033, BF_10_ = 3.035) as well as main effects of group (F_(1,42)_ = 20.78, p < 0.001, ƞ^2^ = 0.251, BF_10_ = 45.545) and RS run (F_(2, 83.85)_ = 192.16, p < 0.001, ƞ^2^ = 0.595, BF_10_ = 3.439*e^28^). Specifically, the greater persistence of task-related hippocampal activity into RS2 in children was absent following the 5-hr consolidation interval. This suggests that the greater post-learning persistence in the hippocampus in children was transient and only observed in the resting state scan immediately following training.

Altogether, children exhibited greater post-learning persistence in the hippocampus but not the other studied regions. This enhanced hippocampal persistence was transient and did not continue throughout the post-consolidation rest epoch.

#### 3.2.3 Persistence x behavior relationships

Separate multiple linear regressions were conducted to assess the relationships between inter-block and post-learning persistence and their respective behavioral measures, as well as age group differences in these associations. Results for exploratory regions are listed in Appendix 4.

Regression models assessing the link between inter-block persistence for each training run and the respective micro-offline performance gains – independent of group – revealed no significant relationships for any primary region of interest in training 1 (HC: b = 0.152, p = 0.631; PUT: b = 0.166, p = 0.741; ACCU: b = 0.105, p = 0.598) or training 2 (HC: b = 0.058, p = 0.835; PUT: b = −0.023, p = 0.961; ACCU: b = 0.215, p = 0.232). There were also no age group differences in the relationships between persistence and micro-offline performance changes (Training Run 1: HC: b = −0.809, p = 0.293; PUT: b = −1.083, p = 0.319; ACCU: b = −0.652, p = 0.270; Training Run 2 HC: b = 0.692, p = 0.331; PUT: b = 0.455, p = 0.660; ACCU: b = −0.083, p = 0.818).

Similarly, post-learning persistence was not significantly related to the behavioral macro-offline performance changes for any region when collapsing across the two age groups (Training 1: HC: b = 0.500, p = 0.154; PUT: b = 0.162, p = 0.692; ACCU: b = 0.117, p = 0.576; Training 2: HC: b = −0.228, p = 0.577; PUT: b = −0.289, p = 0.483; ACCU: b = 0.154, p = 0.441). When assessing age group differences in the associations between post-learning persistence and macro-offline performance changes, no statistically significant differences were revealed for the hippocampus (Tr1: b = 0.560, p = 0.559; Tr2: b = −0.159, p = 0.863) nor the putamen (Tr1: b = 0.503, p = 0.546; Tr2: b = 0.053, p = 0.950) for both training 1 and 2. For the nucleus accumbens control region, no age group differences were present in training 2 (b = 0.418, p = 0.355). There was, however, a trend toward an age group difference for training 1 (b = 0.719, p = 0.093). Note that the difference between age groups was mainly driven by the child participant with an extreme value in the persistence metric. Together, these results provided no evidence for associations between the persistence of task-related activity into offline periods and the magnitudes of the respective offline performance changes.

## 4. Discussion

The current study employed multivoxel correlational structure (MVCS) analyses to investigate whether the developmental advantage in offline consolidation is linked to persistence of learning-related patterns of brain activity into subsequent rest epochs, thought to be reflective of reactivation processes. Our results revealed that: (1) persistence of task activity patterns into inter-block rest was greater in children than adults for the hippocampus, putamen and the exploratory task-relevant regions of interest (i.e., caudate nucleus, primary motor cortex, pre-supplementary motor area, ventral and dorsal premotor cortex, primary sensory cortex); (2) children exhibited greater persistence of task-related activity in the hippocampus, but not in the putamen nor in other exploratory regions, into post-learning rest relative to adults; and, (3) there was no evidence to suggest that the magnitudes of inter-block or post-learning persistence were related to the micro-and macro-offline performance changes, respectively.

### 4.1 Children exhibit greater persistence of task activity patterns into post-learning rest

The current study observed that, consistent with our hypotheses, school-aged children exhibited greater persistence of hippocampal task activity patterns into post-learning rest as compared to adults. This greater persistence observed in children paralleled their enhanced macro-offline motor memory consolidation observed at the behavioral level. Previous research in adults has demonstrated that post-learning hippocampal reactivations reflect the abstract, spatial representation of a learned motor sequence (King et al., 2022). This spatial representation is commonly described as one of the two representations that develop in parallel during MSL, alongside a motoric representation (see Albouy et al., 2013 for a review). Given the enhanced hippocampal persistence in children, one could speculate that they exhibited greater reactivation of this spatial representation relative to adults, potentially resulting in a stronger abstract representation of the motor sequence. This could be interpreted as in line with previous research that suggests a developmental shift from generalization (e.g., regularities/commonalities between events) to specificity (e.g., specific details of events) of memories during childhood, hypothesized to be driven by hippocampal maturation (see Keresztes et al., 2018 for a review). That is, generalizability of the motor memory could be the result of a more abstract representation, as it encodes the motor sequence in allocentric, effector-independent coordinates. In contrast, specificity may be more tightly linked to the rigid, effector-dependent motoric representation.

As the findings from our previous neuroimaging study indicated a link between hippocampal activity and macro-offline performance changes in children (Van Roy et al., 2025), it is tempting to speculate that the enhanced post-learning persistence of hippocampal patterns in children contributes to the developmental advantage in macro-offline consolidation reported here and in previous research (Ashtamker & Karni, 2013; Van Roy et al., 2024). It is worth noting, however, that our regression analyses did not provide evidence for such relationships between post-learning hippocampal persistence and macro-offline performance improvements, including within children. Although we are cautious to not over-interpret the lack of a significant relationship, one potential explanation for the lack of significant brain-behavior associations in the current study could be the difference in the timing of our metrics of interest. Specifically, macro-offline consolidation was measured as the change in performance across the full 5-hour consolidation interval, whereas post-learning persistence was assessed via a single snapshot of brain activity patterns immediately after learning (i.e., during RS2). This limits our ability to make inferences about the neural processes occurring throughout the entire consolidation window. An exploratory follow-up analysis indicated that the greater persistence of hippocampal activity in children was indeed transient, as no group differences were observed at the end of the consolidation interval and immediately prior to the retest (i.e., during RS3). It would be worthwhile for future research to include a denser sampling of post-learning resting state scans to provide a detailed time course of persistence across the 5-hour offline wake interval. A second explanation for the lack of a brain-behavior association is that, and as mentioned above, the greater hippocampal post-learning persistence in children reflects the reactivation of an abstract representation of the motor sequence. As a result, post-learning persistence might be linked to a performance metric that reflects the consolidation of the spatial aspect of the motor sequence rather than a general speed measure. The current task design does not allow us to distinguish between the two representations and future research is certainly warranted.

This is not the first research to speculate about enhanced reactivation processes in children and the potential connections to advantages observed at the behavioral level. Wilhelm et al. (2013) observed greater slow wave activity (SWA) in children, relative to adults, during a night of sleep following implicit learning of a motor sequence task. This SWA was significantly correlated to the enhanced explicit knowledge of the sequence that was observed in children as compared to adults [Wilhelm et al., 2013; but see Voisin et al., 2025 for results reporting similar effects between sleep and wake in children]. As there is evidence indicating that SWA supports an active consolidation process that involves the repeated reactivation of newly encoded memories (Diekelmann & Born, 2010), Wilhelm and colleagues suggested that the developmental advantage in the extraction of explicit sequence knowledge may be the result of more effective SWA-driven reactivation processes. Their finding of a potential childhood benefit linked to post-learning sleep and our observation of enhanced persistence of task-related hippocampal activity during post-learning wake epochs collectively suggest that the offline processing of recently practiced motor skills support developmental advantages in learning and memory-related behaviors.

In contrast to our hypotheses, we found no evidence for differences between children and adults in post-learning persistence of task-related putamen activity. This lies in conflict with a subset of previous literature demonstrating that the putamen still undergoes functional organization (observed in 7-12 year-olds; Greene et al., 2014) as well as structural maturation (age-related changes in 9-17 and 10-30 year-olds; Corrigan et al., 2021; Østby et al., 2009) throughout childhood and into adolescence. Consequently, one could expect that functional activity patterns relating to post-learning reactivation continue to undergo developmental changes as well. As developmental research on the neural mechanisms underlying episodic memory consolidation has largely focused on the hippocampus (e.g., Benear et al., 2022; Ghetti et al., 2010; Sastre et al., 2016), and neuroimaging investigations into motor learning during childhood are relatively limited, the precise role of the putamen in offline consolidation in children remains to be fully elucidated. Nonetheless, the finding of similar persistence in the putamen between age groups would suggest similar offline processing during macro-offline intervals in children and adults.

The patterns of activity observed during post-learning rest were more similar to the task patterns in training run 2 as compared to training run 1 in both age groups. This was observed for both task-relevant regions as well as the control region. One could argue that this training run effect is due to the temporal proximity of training 2 and the post-learning resting-state scan. While we cannot entirely rule out that possibility, the observed results may also reflect differences between training runs that extend beyond mere temporal proximity. For instance, the functional recruitment of the striatum and hippocampus during online task practice has been shown to change as a function of learning (see Albouy et al., 2013 for a review). In this context, patterns of activity observed during training runs 1 and 2 could reflect early and late learning, respectively. The observed persistence into post-learning rest thus may reflect the reactivation of the patterns of brain activity underlying late learning stages in task-relevant regions. However, this interpretation is not fully consistent with previous research in adults that reported greater hippocampal involvement during early learning. Specifically, the magnitude of hippocampal activity is thought to be largest during early learning, which is linked to the acquisition of the hippocampal-mediated abstract representation of the motor sequence (see Albouy et al., 2013 for a review). However, whether the dynamic modulation of the magnitude of the BOLD signal throughout practice relates to multivoxel patterns of activity remains unknown. Moreover, less is known about the dynamical recruitment of the hippocampus during initial learning in children, making it difficult to directly map adult activity patterns onto the developing brain.

### 4.2 Children demonstrate greater inter-block persistence of task-related activity patterns

As hypothesized, children exhibited greater inter-block persistence of hippocampal and putamen task patterns of activity relative to adults, while no such effect was observed in the control region. Similar results were obtained across many of the exploratory, task-relevant ROIs (see Appendix 4), including the caudate nucleus, primary motor and somatosensory cortices, pre-SMA, as well as the ventral and dorsal premotor cortices. This greater inter-block persistence observed in children could be considered in line with the results of the univariate neuroimaging analyses of the same data (Van Roy et al., 2025). Specifically, during initial learning of the sequence, children demonstrated similar levels of activity during active task practice and interleaved rest across a range of brain regions, including the sensorimotor cortex, supplementary motor area and putamen. On the one hand, similar levels of activity could be the result of a particular region *not* being active during active task practice, nor during the inter-block rest modeled as the baseline in the block design. Alternatively, similar levels of activity could be the result of the region being equally active during task practice and the interleaved rest epochs, and perhaps this reflects persistence of task activity into the inter-block rest intervals. Support for the latter interpretation comes from previous research in young adults that linked hippocampal activity during task practice (relative to inter-block rest) to persistence of task-related activity into this offline period (Gann et al., 2023). As both the previously employed univariate and the current multivoxel analyses are inherently limited in their ability to distinguish between underlying mechanisms, greater similarity in activity between task practice and interleaved rest could be a result of either scenario mentioned above. However, it is worth emphasizing that greater inter-block persistence in children was observed across regions known to be recruited during task performance, including the exploratory cortical and subcortical regions, but *not* the nucleus accumbens control region. It could thus be speculated that this pattern of results would likely fit with the explanation that these regions were equally active during blocks of task practice and the interspersed rest.

The greater inter-block persistence of task activity in children may be attributed, at least partially, to the functional organization of the developing brain. Children aged 7-9 years exhibit less functional segregation (Fair et al., 2008, 2009), meaning that their task and rest brain networks are less functionally distinct. This reduced segregation could result in greater overlap (i.e., higher similarity) in the patterns of activity during online task practice and inter-block rest (Thomason et al., 2008), thereby contributing to the higher persistence observed in children. At the same time, greater persistence, particularly during the inter-block intervals, may not solely reflect spontaneous offline reactivation processes. Although offline processing of recently learned information is traditionally considered an “unconscious”, spontaneous process, Camacho et al. (2020) proposed that the developing brain is actively and consciously engaged during rest periods. Accordingly, the greater inter-block persistence observed in children could arise from active rehearsal of the motor sequence. As the methodology employed here cannot distinguish between these possibilities, it remains unclear whether the greater persistence in children reflects spontaneous reactivation or active rehearsal processes.

In contrast to our hypothesis, we found no evidence to suggest that inter-block persistence was related to the behavioral marker of micro-offline consolidation within or across age groups for any brain region examined. This is inconsistent with the association between hippocampal replay and micro-offline performance changes that was previously reported in adults (Buch et al., 2021). It is worth nothing that Buch et al. (2021) employed magnetoencephalography (MEG), enabling them to extract replay metrics that are fundamentally different than the current ones from a temporal point of view. This methodological difference could explain the inconsistency in findings. In fact, a recent study that employed similar multivariate fMRI analyses as in the current study also did not observe an association between micro-offline reactivation and performance changes in adults (Gann et al., 2023). Taken together, these findings raise the possibility that enhanced micro-offline reactivation in children may contribute to their behavioral advantage; however, such a brain-behavior correlation may emerge only when the temporal features of replay can be extracted.

### 4.4 Interblock versus post-learning persistence

Children exhibited greater inter-block persistence than adults across most task-relevant brain regions examined, whereas their enhanced post-learning persistence was observed only in the hippocampus. A similar pattern emerged in our previous neuroimaging study (Van Roy et al., 2025), in which children showed smaller modulations of brain activity between task and interleaved rest periods across numerous regions – consistent with continued engagement of these task-relevant regions during inter-block rest - but age group differences across the macro-offline period were limited to a smaller set, including the hippocampus, primary motor cortex and cerebellum. These results collectively suggest that developmental differences in the offline processing of recently practiced motor skills *differ* between the two timescales (i.e., micro- and macro-offline). As we did not observe any significant associations between offline performance improvements and offline persistence metrics, the functional significance of these differences is not yet known and warrants attention in future research.

### 4.5 Limitations

There are several limitations and methodological considerations of the current research that warrant further attention. The following section focuses on limitations specific to the multivoxel neuroimaging approach. Considerations related to experimental procedures and behavioral analyses are outlined in the discussion of Van Roy et al. (2025). *First*, and mentioned previously, the low temporal resolution of fMRI and the implementation of the motor learning task in the current study do not afford the extraction of temporal characteristics of post-learning reactivation. Conflicting findings in prior literature indicate that this inability to distinguish between temporal features (e.g., sequence-specific patterns of activity) may have masked associations between offline persistence and the respective behavioral performance changes, at least on the micro-timescale (Buch et al., 2021; Gann et al., 2023). Whereas no prior study has examined the link between post-learning reactivation and macro-offline performance changes in MSL, correlations between replay and behavior have been shown in episodic research that employed fMRI techniques (Schapiro et al., 2018; Tambini & Davachi, 2013). Future research is warranted to disentangle the relationship between specific features of post-learning reactivation and offline performance changes on a motor sequence task.

*Second*, one could argue that the detrimental influence of head motion artifacts is magnified in studies that employ a multivoxel analytical approach due to its spatial sensitivity. For instance, structured motion-related noise across voxels could lead to correlations that do not reflect neural activity per se, whereas motion-related variability as well as voxel misalignment can disrupt real neural correlations. Furthermore, reduced signal-to-noise ratio can introduce individual variability and thus, make the detection of group differences harder. This deleterious effect is likely amplified in a pediatric sample, as children tend to move more during scanning sessions as compared to adults (see Appendix 3). Although we adopted multiple approaches to minimize the impact of head movement (e.g., exclusion of participants with a maximum linear movement > 5mm, disregarding high-motion volumes with FD > 0.5, and implementing head motion parameters as nuisance regressors), we cannot fully eliminate the possibility that head motion – and differences in the magnitude of head motion between children and adults - biased the MVCS results.

*Third*, MVCS approaches similar to the one employed here adopt a within-subject control condition to assess the magnitude of persistence. For example, our macro-offline persistence analysis compared the similarity between online task practice and post-learning rest (RS2) with the similarity between the same task practice and pre-learning rest. We implemented the same strategy for micro-offline persistence. However, the use of pre-learning rest as the within-subject control may not be an ideal control given differences with the inter-block rest periods. Specifically, participants saw a dark screen with a white cross during the resting-state scans, whereas visual input during interleaved rest consisted of a red cross below a fixed sequence of symbols. Furthermore, while the acquired data was continuous – except for scrubbed volumes - during pre-learning rest, inter-block MVCS matrices were obtained by extracting and concatenating the images during the inter-block rest periods. Similarly, the computation of MVCS matrices for online task practice also included such extraction and concatenation procedures. This likely resulted in greater similarity in activity patterns between online task practice and inter-block rest and smaller similarity between task practice and pre-learning rest, resulting in the robust increases we observed in similarity with task practice from RS1 to inter-block rest. Due to these aforementioned differences, we were not able to assess whether there was significant micro-offline persistence *per se* (i.e., whether the similarity between task and inter-block rest was significantly greater than task and pre-learning rest). Future research could use the interleaved rest breaks of a similar task that does not include learning (e.g., random serial reaction time task) as a within-subject control for inter-block persistence. Nevertheless, these epoch-based differences are expected to be similar between age groups and thus, cannot fully account for the greater micro-offline persistence that was observed for children.

*Fourth,* the nucleus accumbens served as a control region, based on its minimal role in motor memory and its subcortical location providing a signal-to-noise ratio comparable to the hippocampus and putamen. However, given the small size of the nucleus accumbens, its signal may be more impacted by noise due to partial-volume effects and head movement than larger regions such as the putamen. Nevertheless, this region appeared to be the best choice, as other subcortical regions could be considered as task-relevant regions (e.g., globus pallidus).

*Fifth,* post-learning persistence was assessed using a 6-minute post-learning resting-state scan, providing only a snapshot into reactivation processes during the macro-offline consolidation period. This limits our ability to make inferences about neural processes occurring throughout the entire consolidation period. Furthermore, and as discussed, it may have obscured potential brain-behavior relationships. While the inclusion of a post-consolidation resting-state scan allowed us to examine how activity patterns changed from the beginning to the end of the consolidation interval, future studies should consider including resting-state imaging at multiple timepoints throughout this offline period.

### 4.5 Conclusions

The current study employed a multivoxel neuroimaging approach to assess the persistence of learning-related patterns of neural activity into post-task rest epochs in children and adults. Our results revealed greater persistence of learning-related activity patterns in the hippocampus, putamen and exploratory regions during the short rest periods that are commonly interspersed between blocks of task practice in children as compared to adults. Children also exhibited greater persistence of task-related hippocampal activity patterns into post-learning rest. These results suggest that the childhood advantage in motor memory consolidation across periods of wakefulness may be supported, at least partially, by an enhanced reactivation of task-dependent activity patterns during these offline epochs. Future research is warranted to unravel characteristics of the enhanced post-learning persistence of learning-related patterns of brain activity in children as well as the link to behavioral consolidation measures.

