## Supplemental material for "Children exhibit greater persistence of motor learning-related patterns of hippocampal activity into post-task wake epochs"

**Supplementary information**

**Authors**

Anke Van Roy[1], Ainsley Temudo[1], Emily K. Taylor[1], Vincent Koppelmans[2,3], Kerstin Hödlmoser[4,5], Genevieve Albouy[1], Bradley R. King[1*]

**Affiliations**

[1] Department of Health and Kinesiology, College of Health, University of Utah, Salt Lake City, UT 84112, USA

[2] Department of Psychiatry, University of Utah, Salt Lake City, UT 84112, USA

[3] Huntsman Mental Health Institute, University of Utah, Salt Lake City, UT 84112, USA

[4] Department of Psychology, University of Salzburg, 5020 Salzburg, Austria

[5] Centre for Cognitive Neuroscience Salzburg, University of Salzburg, 5020 Salzburg, Austria

[*] **Corresponding author**

Bradley King, PhD
Department of Health & Kinesiology, College of Health, University of Utah
250 S 1850 E
Salt Lake City, Utah 84112


**Table S11.** Statistics of linear regression analyses assessing the relationships between post-learning persistence and macro-offline performance changes in exploratory brain regions. 27

**Figure S8**. Relationships between macro-offline performance changes and post-learning persistence for the ventral premotor cortex (PMC) for children (blue) and adults (orange). 28

**Table S12.** Results of group x RS ANOVAs comparing persistence into the post-consolidation RS3 vs. persistence into RS2. 29

**Figure S9.** Persistence of task activity during training 2 into post-learning (i.e., RS2) and post-consolidation rest (i.e., RS3). 30

**Figure S10.** Similarity of task activity patterns during training 1 (left) and training 2 (right) with patterns of activity during pre-learning (i.e., RS1), post-learning (i.e., RS2) and post-consolidation rest (i.e., RS3). Similarity indices were computed using r-to-z transformed correlations. 31

**Supplementary references** 34

### APPENDIX 1: Protocol deviations

**Table S1.** Registration deviations.

List of the deviations from the registered analyses followed by their justification. Deviations are marked with an asterisk in the main text.

| | Registered | Final report |
|---|---|---|
| 1 | Spatial pre-processing will be performed in **SPM12 in Matlab**. The structural T1-weighted image is reoriented to the Montreal Neurological Institute (MNI) template. The reoriented image is then segmented into gray matter, white matter, cerebrospinal fluid (CSF), bone, soft tissue, and background. Task-based and resting-state functional volumes of each participant are realigned to the first image of each MRI run and subsequently, realigned to the mean functional image computed across MRI runs of the respective participant using rigid body transformations. The mean image is co-registered to the pre-processed high-resolution T1-weighted structural image using a rigid body transformation optimized to maximize the normalized mutual information between the two images. The resulting co-registration parameters are then applied to the realigned functional images. | Initial spatial preprocessing was performed in **fMRIPrep 24.1.0**. The T1w anatomical image was corrected for intensity non-uniformity, skull-stripped and segmented into white matter (WM), gray matter (GM) and cerebrospinal fluid (CSF). For each task-related and resting-state run, head motion was estimated (see Supplemental Tables S3 and S4 in Appendix 3 for information on the magnitude of head motion parameters and statistical comparisons, respectively), **and functional images were corrected for susceptibility distortion using GRE fieldmaps as well as co-registered to anatomical space.** Several confounding time-series were calculated based on the preprocessed BOLD data, such as framewise displacement (FD), and anatomical noise components (i.e., CSF and WM probabilistic maps) were identified using principal component analysis (PCA; explaining >50 % of variance; see below for details on preprocessing steps that used these confounding time-series). Consistent with previous research employing similar multivoxel analytic approaches (King et al., 2022; Tambini & Davachi, 2013), anatomical and functional images remained in native space (i.e., not normalized to a template). |
| | **Justification**: Pre-processing was performed in fMRIPrep 24.1.0 instead of SPM12 to enable the implementation of correction for inhomogeneities in the magnetic field. Although SPM12 provides this option as well, prior researchers in our team experienced unforeseen image distortions when implementing fieldmaps using SPM. | |
| 2 | If the group x training run ANOVA assessing age groups in the similarity between activity patterns during online task practice of each training run and *pre-learning* rest (i.e., RS1) produces a significant main effect of age group and/or an age group by epoch | Age group differences in the similarity with pre-learning rest (RS1) were accounted for by **computing a normalized persistence measure** by subtracting the SI between RS1 and online task practice from the SI between online task practice and the rest epoch of |

| | | |
|---|---|---|
| | interaction, our confirmatory analyses will be modified to account for this. Specifically, **RS1 will be included into the statistical models**, resulting in group (children vs. adults) x rest epoch (RS1 vs. rest epoch of interest) ANOVAs. **Separate statistical models will be conducted per training run**. | interest (i.e., SI (online practice \| rest epoch of interest) – SI (RS1 \| online practice)). These normalized persistence scores were then subjected to mixed ANOVAs with group as the between-subject factor and **training run as the within-subject factor.** |
| | **Justification:** We elected to create a new measure that indicates the relative difference between similarities with pre- and post-task rest rather than statistically comparing them given the notable differences in the nature of the signal between RS1 and the interleaved rest periods, such as the visual stimuli (i.e., dark screen with white cross vs. fixed sequence of symbols in red) and how the signals for the MVCS matrices were obtained (i.e., extracted and concatenated for interleaved rest vs. full run for RS1). The same approach was implemented for the assessment of post-learning persistence. | |

**Table S2.** Deviations from the standard protocol.

There were occasional issues that arose during data acquisition, particularly in young children, that resulted in deviations from the standard protocol.

| Subject | Age (yrs) | Deviation |
|---|---|---|
| E4 | 7.92 | There was excessive head motion during the inter-block rest periods of training runs 1 and 2. Specifically, the removal of volumes with excessive head movement (maximum translation > 5 mm and FD > 0.5 mm) resulted in the exclusion of more than 50% of volumes during interleaved rest. Accordingly, this participant had missing data for the assessment of inter-block persistence for both training runs 1 and 2. |
| E31 | 9.58 | The participant was no longer following task instructions during the post-learning test (i.e., to perform the task as accurately and quickly as possible), indicated by an accuracy of less than 40% and a decrease in the time between transitions of over 150% percent. As this behavior was only evident during the post-learning test phase but not the training runs that were used in the fMRI analyses, we elected to include this participant in analyses, but the computation of their macro-offline performance changes was altered. Specifically, this metric was computed as the change in performance from the last two blocks of training 2 (in contrast to the average of the two test blocks) to the first two blocks of retest. Note that this resulted in a substantially smaller macro-offline gain and thus was in the direction opposite to our hypothesized (and reported) results.<br>If we maintained the original computation (i.e., based on the average of the two test blocks), the significant group differences in macro-offline changes in performance speed and accuracy were as follows: Speed: $t_{43} = 2.602$, $p = 0.006$, $G = 0.76$, $BF_{10} = 4.085$; accuracy: $t_{43} = 2.311$, $p = 0.015$, $G = 0.67$, $BF_{10} = 2.580$.<br><br>Due to excessive head motion (i.e., maximum translation exceeding 5 mm), volumes 1 to 82 of Training 2 were excluded from the MVCS analysis. These volumes included the initial rest period as well as active task practice of block 1. |
| E45 | 8.17 | There was excessive head motion during task practice and inter-block rest periods of training run 2. Specifically, the removal of volumes with excessive head movement (maximum translation > 5 mm and FD > 0.5 mm) resulted in the exclusion of more than 50% of volumes during both task and rest blocks. Accordingly, this participant had missing data for the assessments of post-learning persistence and inter-block persistence for training run 2. |
| E46 | 25.33 | Due to difficulties with data reconstruction, RS2 includes 229 images instead of 375. |
| E49 | 10 | Due to excessive head motion (i.e., maximum translation exceeding 5 mm), volumes 284-375 of RS1 as well as volumes 719-914 of Training 2 were excluded from the MVCS analysis. |

| | | |
|---|---|---|
| E54 | 11.08 | Due to difficulties with data reconstruction RS2 only includes 230 images instead of 375. |
| E69 | 18.92 | Due to difficulties with data reconstruction, RS2 includes 229 images instead of 375. |
| E75 | 18.17 | Due to difficulties with data reconstruction, RS2 includes 304 images instead of 375. |
| E96 | 24.08 | RS2 included a corrupted volume. As a result, this resting-state run included 374 images instead of 375. |
| E101 | 8.25 | Due to excessive head motion (i.e., maximum translation exceeding 5 mm), volumes 250-375 of RS1 were excluded from the MVCS analysis. |
| E103 | 10.83 | Participant lost their finger placement on the keyboard at the beginning of block 6 of training run 1. Scanning was stopped to reposition the fingers and the participant completed 5 additional blocks of training run 1. Functional images and performance during this portion of block 6 of the first part of training 1 were ignored. |
| E107 | 8.67 | There was excessive head motion during task practice and the inter-block rest periods of training runs 1 and 2. Specifically, the removal of volumes with excessive head movement (maximum translation > 5 mm and FD > 0.5 mm) resulted in the exclusion of more than 50% of volumes during both task and rest blocks of both training runs. Accordingly, this participant had missing data for the assessments of post-learning persistence and inter-block persistence for both training runs. |
| E113 | 7.25 | Due to excessive head motion (i.e., maximum translation exceeding 5 mm), volumes 1008 to 1293 of task practice during training 2 were excluded from the post-learning MVCS analysis.<br><br>There was excessive head motion during inter-block rest periods of training run 1. Specifically, the removal of volumes with excessive head movement (maximum translation > 5 mm and FD > 0.5 mm) resulted in the exclusion of more than 50% of volumes during rest blocks of this training run. Accordingly, this participant had missing data for the assessment of inter-block persistence for training run 1. |
| E138 | 7.58 | There was excessive head motion during inter-block rest periods of training run 1. Specifically, the removal of volumes with excessive head movement (maximum translation > 5 mm and FD > 0.5 mm) resulted in the exclusion of more than 50% of volumes during rest blocks of this training run. Accordingly, this participant had missing data for the assessment of inter-block persistence for training run 1. |

#### APPENDIX 2: Supplementary methods

Initial spatial preprocessing was performed in fMRIPrep 24.1.0 (Esteban et al., 2019). The text below was automatically generated by fMRIPrep in each participant's visual report and is released under the CC0 license. Additions or clarifications to this automated text made by the authors of this manuscript are explicitly noted below.

##### 2.1 fMRIPrep pre-processing details

A total of 2 T1-weighted (T1w) images were found within the input BIDS dataset [Author note: the acquired T1w image was duplicated, as fmriprep requires a T1w image per testing session directory (i.e., training and retest sessions)]. Each T1w image was corrected for intensity non-uniformity (INU) with N4BiasFieldCorrection (Tustison et al., 2010), distributed with ANTs 2.5.3 (Avants et al., 2008; RRID:SCR_004757). The T1w-reference was then skull-stripped with a Nipype implementation of the antsBrainExtraction.sh workflow (from ANTs), using OASIS30ANTs as target template. Brain tissue segmentation of cerebrospinal fluid (CSF), white-matter (WM) and gray-matter (GM) was performed on the brain-extracted T1w using fast (FSL (version unknown), RRID:SCR_002823, Zhang et al., 2001). An anatomical T1w-reference map was computed after registration of 2 <module 'nipype.interfaces.image' from '/opt/conda/envs/fmriprep/lib/python3.11/site-packages/nipype/interfaces/image.py'> images (after INU-correction) using mri_robust_template (FreeSurfer 7.3.2, Reuter et al., 2010). Volume-based spatial normalization to one standard space (MNI152NLin2009cAsym) was performed through nonlinear registration with antsRegistration (ANTs 2.5.3), using brain-extracted versions of both T1w reference and the T1w template. The following template was were selected for spatial normalization and accessed with TemplateFlow (24.2.0, Ciric et al., 2022): ICBM 152 Nonlinear Asymmetrical template version 2009c [Fonov et al., 2009, RRID:SCR_008796; TemplateFlow

ID: MNI152NLin2009cAsym][Author note: The spatial normalization to MNI space described above was not used for the current analyses. Rather, voxel-based analyses require images to be in native space].

For each of the 7 BOLD runs found per subject (across all tasks and sessions) [Author note: the post-learning test was included in the fMRIPrep processing as it was part of the BIDS structured data], the following preprocessing was performed. First, a reference volume was generated, using a custom methodology of fMRIPrep, for use in head motion correction. Head-motion parameters with respect to the BOLD reference (transformation matrices, and six corresponding rotation and translation parameters) were estimated before any spatiotemporal filtering using mcflirt (FSL , Jenkinson et al., 2002) [Author note: Information on the magnitude of head motion parameters, such as FD, translation and rotation, is provided in Tables S3 and S4]. The estimated fieldmap was then aligned with rigid-registration to the target EPI (echo-planar imaging) reference run. The field coefficients were mapped on to the reference EPI using the transform. The BOLD reference was then co-registered to the T1w reference using mri_coreg (FreeSurfer) followed by flirt (FSL, Jenkinson & Smith, 2001) with the boundary-based registration (Greve & Fischl, 2009) cost-function. Co-registration was configured with six degrees of freedom. Several confounding time-series were calculated based on the preprocessed BOLD: framewise displacement (FD), DVARS and three region-wise global signals. FD was computed using two formulations following Power (absolute sum of relative motions, Power et al., 2014) and Jenkinson (relative root mean square displacement between affines, Jenkinson et al., 2002). FD and DVARS were calculated for each functional run, both using their implementations in Nipype (following the definitions by Power et al., 2014). The three global signals were extracted within the CSF, the WM, and the whole-brain masks. Additionally, a set of physiological regressors were extracted to allow for component-based noise correction (CompCor, Behzadi et al., 2007). Principal components were estimated after high-pass filtering the preprocessed BOLD time-series (using a discrete cosine filter with 128s cut-off) for the two CompCor variants: temporal (tCompCor) and anatomical (aCompCor). tCompCor components were then calculated from the top 2% variable voxels within the brain mask. For aCompCor, three probabilistic masks (CSF, WM and combined CSF+WM) were generated in anatomical space. The implementation differs from that of Behzadi et al. in that instead of eroding the masks by 2 pixels on BOLD space, a mask of pixels that likely contain a volume fraction of GM is subtracted from the aCompCor masks. This mask is obtained by thresholding the corresponding partial volume map at 0.05, and it ensures components are not extracted from voxels containing a minimal fraction of GM. Finally, these masks were resampled into BOLD space and binarized by thresholding at 0.99 (as in the original implementation). Components were also calculated separately within the WM and CSF masks. For each CompCor decomposition, the k components with the largest singular values were retained, such that the retained components' time series are sufficient to explain 50 percent of variance across the nuisance mask (CSF, WM, combined, or temporal). The remaining components were dropped from consideration. The head-motion estimates calculated in the correction step were also placed within the corresponding confounds file. The confound time series derived from head motion estimates and global signals were expanded with the inclusion of temporal derivatives and quadratic terms for each (Satterthwaite et al., 2013). Frames that exceeded a threshold of 0.5 mm FD or 1.5 standardized DVARS were annotated as motion outliers. Additional nuisance timeseries were calculated by means of principal components analysis of the signal found within a thin band (crown) of voxels around the edge of the brain, as proposed by (Patriat, Reynolds, and Birn 2017).

All resamplings can be performed with a single interpolation step by composing all the pertinent transformations (i.e. head-motion transform matrices, susceptibility distortion correction when available, and co-registrations to anatomical and output spaces). Gridded (volumetric) resamplings were performed using nitransforms, configured with cubic B-spline interpolation.

**2.2 Inclusion of pre-learning RS1**

Prior to assessing task-related persistence into post-learning offline epochs, and consistent with our registration, we first examined whether the age groups differed in the similarity between activity patterns during online task practice of each training run and pre-learning rest (i.e., RS1). Specifically, SIs were computed between RS1 and online task practice in training run 1 (i.e., fMRI images obtained during online task practice from training 1 were extracted and concatenated) as well as training run 2 (extracted and concatenated). The SIs between RS1 and online task practice of each training run were then subjected to a mixed ANOVA with group (2 levels: children versus adults) as between-subject factor and run (2 levels: training 1 versus training 2) as within-subject factor. We registered that if analyses did *not* reveal a significant main effect of age group and/or a group x epoch interaction, the subsequent analyses would exclude RS1 as an epoch of interest and simply assess between-group differences in the SIs between online task practice and the rest epoch of interest. However, the pre-analysis check found group effects for the hippocampus and the nucleus accumbens control region (hippocampus: $F_{(1,42)} = 10.12$, $p = 0.003$, $\eta^2 = 0.168$, $BF_{10} = 14.025$; putamen: $F_{(1,42)} = 0.20$, $p = 0.749$, $\eta^2 = 0.002$, $BF_{10} = 0.460$; nucleus accumbens $F_{(1,42)} = 7.68$, $p = 0.008$, $\eta^2 = 0.146$, $BF_{10} = 5.137$), with higher similarity indices in children as compared to adults. Consequently, age group differences in the similarity in activity patterns between online task practice and baseline rest were accounted for in the further analyses of all studied regions*.

#### APPENDIX 3: Head motion and voxels included

**Table S3.** Head motion parameters, as well as the number of volumes included in analyses and percentage of excluded volumes following scrubbing procedures.

| **Epoch/region of interest** | | **Children** | **Adults** |
|---|---|---|---|
| *A. Motion parameters* | | | |
| RS1 | FD (avg.) | 0.166 [0.105-0.220] | 0.116 [0.079-0.165] |
| | Translation (max.) | 0.882 [0.362-2.625] | 0.719 [0.113-1.257] |
| | Rotation (max.) | 0.014 [0.003-0.064] | 0.006 [0.001-0.025] |
| Training 1 | FD (avg.) | 0.200 [0.134-0.277] | 0.141 [0.082-0.209] |
| | Translation (max) | 1.495 [0.424-3.128] | 0.944 [0.177-3.226] |
| | Rotation (max.) | 0.034 [0.003-0.195] | 0.009 [0.001-0.026] |
| Training 2* | FD (avg.) | 0.193 [0.121-0.273] | 0.132 [0.083-0.187] |
| | Translation (max) | 1.141 [0.341-2.407] | 0.684 [0.158-2.572] |
| | Rotation (max.) | 0.025 [0.004-0.073] | 0.006 [0.001-0.023] |
| RS2 | FD (avg.) | 0.154 [0.092-0.216] | 0.102 [0.063-0.152] |
| | Translation (max) | 1.020 [0.132-2.877] | 0.374 [0.097-1.134] |
| | Rotation (max.) | 0.013 [0.001-0.084] | 0.004 [0.0003-0.018] |
| *B. Included volumes* | | | |
| RS1 | | 333 [226 – 375] | 373 [362 – 375] |
| Training 1 | Task blocks | 540 [269 – 869] | 285 [136 – 470] |
| | Rest periods$ | 217 [153 – 273] | 245 [199 – 261] |
| Training 2 | Task blocks* | 409 [116 – 785] | 206 [102 – 382] |
| | Rest periods# | 206 [134 – 267] | 247 [216 – 263] |
| RS2 | | 324 [224 – 375] | 355 [229 – 375] |
| *C. Percentage of excluded volumes* | | | |
| RS1 | | 11 [0 – 40] | 1 [0 – 3] |
| Training 1 | Task blocks | 18 [0 – 58] | 3 [0 – 17] |
| | Rest periods$ | 18 [0 – 45] | 3 [0 – 21] |
| Training 2 | Task blocks* | 18 [0 – 68] | 3 [0 – 17] |
| | Rest periods# | 21 [0 – 47] | 2 [0 – 13] |
| RS2 | | 12 [0 – 35] | 1 [ 0 – 7] |

Section A shows head motion parameters, including average framewise displacement (FD), maximum absolute values of the translation and rotation parameters per age group for each run. Translation and rotation maxima were computed across all three axes (x, y and z). Sections B and C display the average number of brain volumes included as well as the mean percentage of volumes excluded, respectively, following the scrubbing procedure based on FD >0.5 and translation >5mm. Minima and maxima across participants are shown between brackets. Values for adults are always based on 23 datasets. Values for children are based on 21 datasets unless indicated otherwise. * = 20 children; # = 19 children; $ = 18 children.

**Table S4.** Statistics of group x run ANOVAs on head motion parameters.

| Variable | df | F | p | $\eta^2$ | $BF_{10}$ |
|---|---|---|---|---|---|
| *A. Average framewise displacement* | | | | | |
| Group | 1, 41 | 41.06 | < 0.001* | 0.425 | $2.37^{*}e^{5}$ |
| Run | 2.13, 87.15 | 43.81 | < 0.001* | 0.219 | $5.499^{*}e^{16}$ |
| Group x Run | 2.13, 87.15 | 0.85 | 0.437 | 0.005 | 0.168 |
| *B. Maximum translation* | | | | | |
| Group | 1, 41 | 14.76 | < 0.001* | 0.119 | 59.168 |
| Run | 2.28, 93.46 | 6.03 | 0.002* | 0.084 | 130.575 |
| Group x Run | 2.28, 93.46 | 1.33 | 0.269 | 0.020 | 0.329 |
| *C. Maximum rotation* | | | | | |
| Group | 1, 41 | 23.85 | < 0.001* | 0.143 | 230.867 |
| Run | 1.70, 69.59 | 3.97 | 0.029* | 0.065 | 2.857 |
| Group x Run | 1.70, 69.59 | 2.14 | 0.133 | 0.036 | 0.791 |

Statistical output of the analyses assessing effects of group (children vs. adults) and run (RS1, training 1, training 2, RS2) on head motion parameters (see Table S3 above for means and ranges). Significant values are marked with an asterisk. n = 20 children, 23 adults. As expected, children exhibited significantly more head motion across all three motion variables as compared to adults.

**Table S5.** Number of voxels included in the MVCS matrices per studied region.

| Epoch/region of interest | Children | Adults |
|---|---|---|
| Hippocampus | 692 [600 – 835] | 728 [612 – 826] |
| Putamen | 698 [459 – 859] | 539 [428 – 663] |
| Nucleus accumbens | 130 [87 – 172] | 124 [94 – 166] |
| Caudate nucleus | 689 [548 – 852] | 660 [488 – 907] |
| Primary motor cortex | 2224 [1690 – 2656] | 2123 [969 – 2700] |
| Pre-supplementary motor area | 594 [450 – 891] | 589 [231 – 856] |
| Supplementary motor area | 728 [363 – 987] | 725 [438 – 1054] |
| Ventral premotor area | 2094 [1593 – 2580] | 1958 [1108 – 2541] |
| Dorsal premotor area | 1594 [1070 – 2217] | 1451 [476 – 2169] |
| Primary somatosensory cortex | 1483 [1204 – 1756] | 1439 [667 – 1803] |

Values represent the average number of voxels included, with ranges in between brackets. Hippocampus and putamen were the primary regions of interest. The nucleus accumbens served as a control. All other regions of interest were considered exploratory (i.e., not included in the registration), with results presented in this supplemental material.

#### APPENDIX 4: Supplemental MVCS results

##### 4.1. Persistence into inter-block rest periods

**Table S6.** Results of regressions that tested the relationship between age and inter-block persistence within the sample of children.

| | Inter-block Persistence | |
|---|---|---|
| **Region of interest** | **Training 1** | **Training 2** |
| Hippocampus | $R^2$ = 0.026; p = 0.645 | $R^2$ = 0.017; p = 0.405 |
| Putamen | $R^2$ = 0.206; p = 0.026* | $R^2$ = 0.002; p = 0.524 |
| Nucleus accumbens | $R^2$ = 0.465; p = 0.308 | $R^2$ = 0.199; p = 0.283 |
| Caudate nucleus | $R^2$ = 0.107; p = 0.127 | $R^2$ = 0.008; p = 0.672 |
| Primary motor cortex | $R^2$ = 0.001; p = 0.889 | $R^2$ = 0.010; p = 0.669 |
| Pre-supplementary motor area | $R^2$ = 0.003; p = 0.822 | $R^2$ = 0.002; p = 0.864 |
| Supplementary motor area | $R^2$ = 0.033; p = 0.433 | $R^2$ = 0.065; p = 0.263 |
| Ventral premotor area | $R^2$ = 0.005; p = 0.767 | $R^2$ = 0.014; p = 0.610 |
| Dorsal premotor area | $R^2$ = 0.026; p = 0.483 | $R^2$ = 0.126; p = 0.114 |
| Primary somatosensory cortex | $R^2$ = 0.017; p = 0.570 | $R^2$ = 0.016; p = 0.590 |

$R^2$ = R-squared. * indicates significant relationships (uncorrected for multiple comparisons). Results are visualized in Figures S1-S2 below.

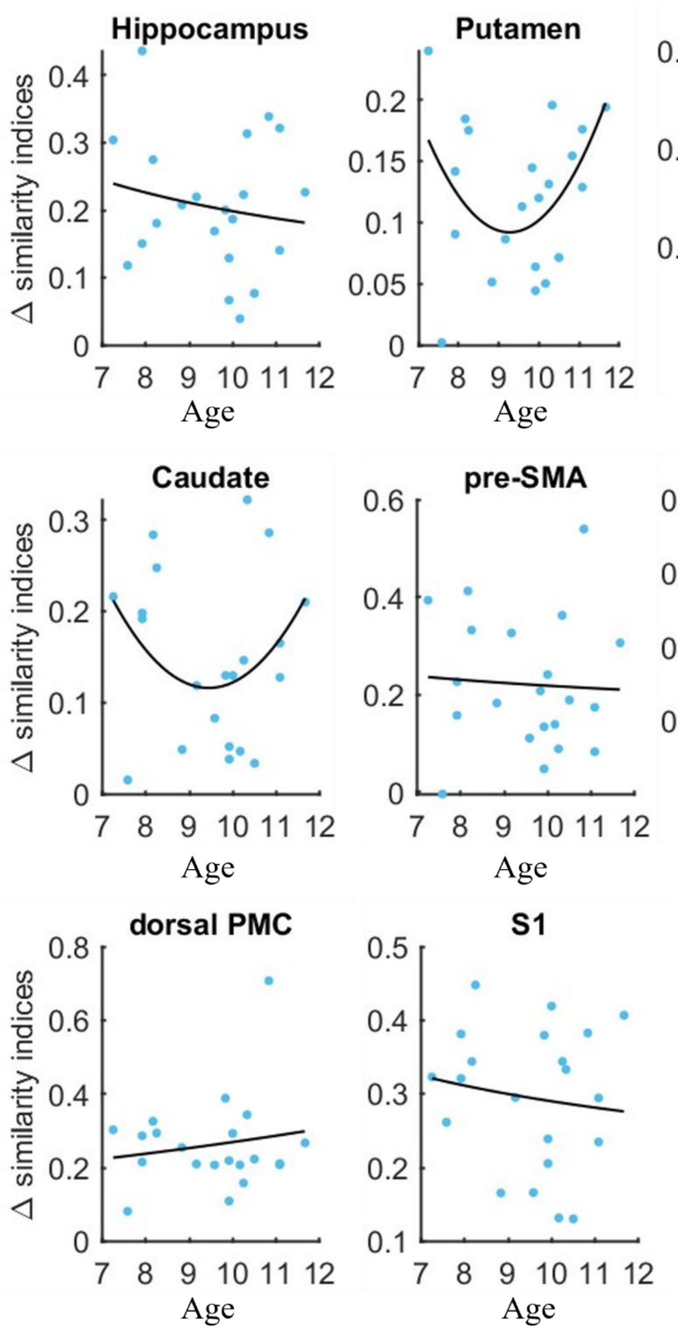

**Figure S1.** Inter-block persistence during training 1 plotted as a function of age within children for primary and exploratory regions. Potential fit options were single exponential, double exponential, linear, quadratic and power functions. M1 = primary motor cortex, pre-SMA = pre-supplementary motor area, SMA = supplementary motor area, PMC = premotor cortex, S1 = primary somatosensory cortex. Statistics are provided in Supplementary Table S6. The relationship between age and inter-block persistence in the putamen was statistically significant ($p = 0.026$, uncorrected for multiple comparisons). The relationship was best fit with a quadratic function, with greater persistence observed in the younger (~7 years) and older children (~11 years).

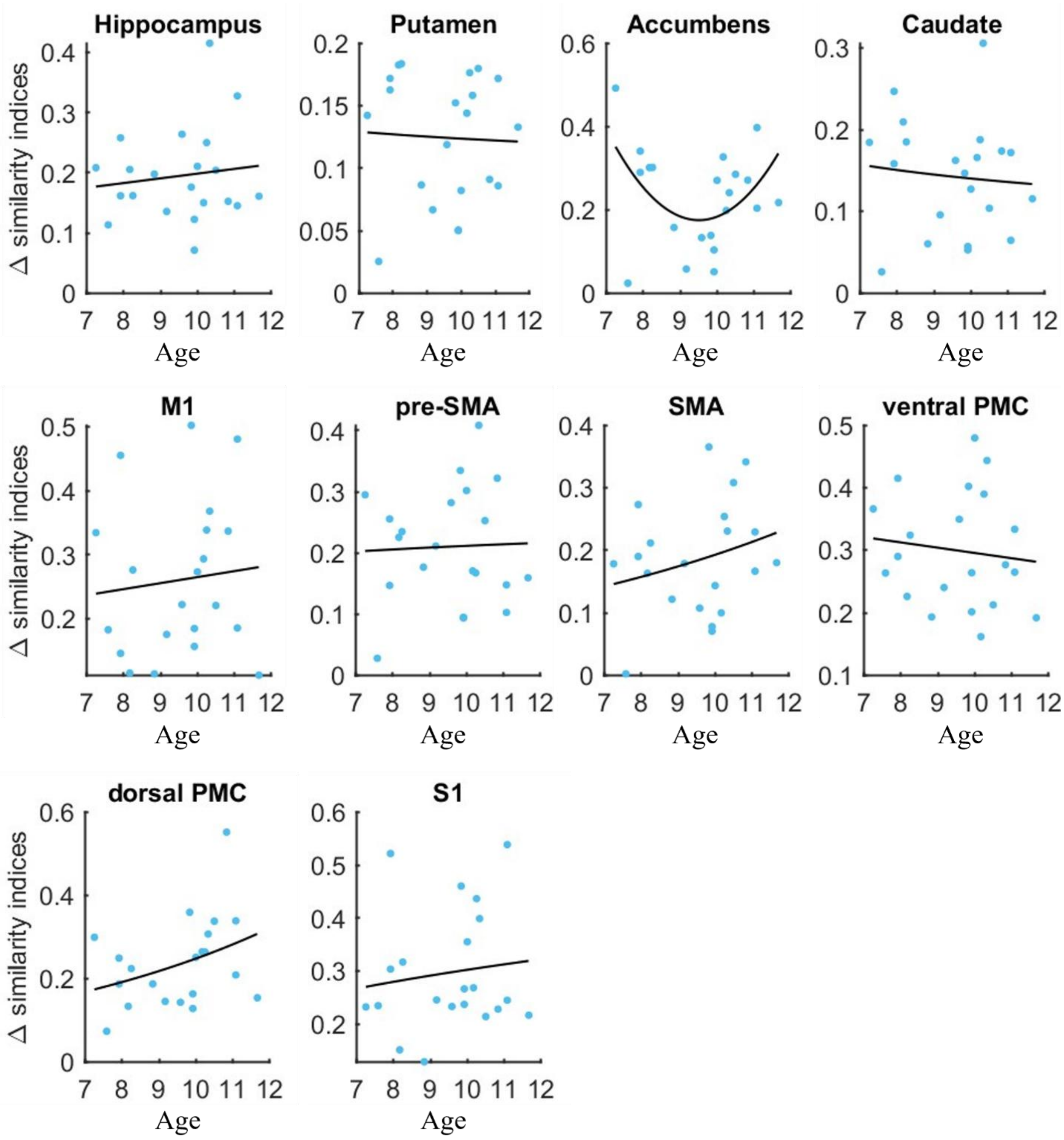

**Figure S2.** Inter-block persistence during training 2 plotted as a function of age within children for primary and exploratory regions. Potential fit options were single exponential, double exponential, linear, quadratic and power functions. M1 = primary motor cortex, pre-SMA = pre-supplementary motor area, SMA = supplementary motor area, PMC = premotor cortex, S1 = primary somatosensory cortex. Statistics are provided in Supplementary Table S6. None of the depicted relationships were significant at $p<0.05$.

**Table S7.** Results of group x run ANOVAs on inter-block persistence in exploratory regions.

| Region | Effect | df | F | p | $\eta^2$ | $BF_{10}$ |
|---|---|---|---|---|---|---|
| Caudate nucleus | Group | 1, 38 | 17.14 | < 0.001* | 0.270 | 137.58 |
| | Training run | 1, 38 | 0.14 | 0.707 | < 0.001 | 0.239 |
| | Group x run | 1, 38 | 0.52 | 0.474 | 0.002 | 0.495 |
| Primary motor cortex | Group | 1, 38 | 4.85 | 0.034* | 0.098 | 3.121 |
| | Training run | 1, 38 | 0.85 | 0.361 | 0.003 | 0.357 |
| | Group x run | 1, 38 | 0.50 | 0.482 | 0.002 | 0.372 |
| Pre-supplementary motor area | Group | 1, 38 | 5.78 | 0.021* | 0.116 | 2.886 |
| | Training run | 1, 38 | 0.19 | 0.663 | < 0.001 | 0.262 |
| | Group x run | 1, 38 | 0.26 | 0.610 | < 0.001 | 0.304 |
| Supplementary motor area | Group | 1, 38 | 4.02 | $0.052^{+}$ | 0.083 | 1.511 |
| | Training run | 1, 38 | 0.16 | 0.688 | < 0.001 | 0.260 |
| | Group x run | 1, 38 | 0.12 | 0.731 | < 0.002 | 0.501 |
| Ventral premotor area | Group | 1, 38 | 28.15 | < 0.001* | 0.382 | 2785.02 |
| | Training run | 1, 38 | 1.38 | 0.247 | 0.006 | 0.474 |
| | Group x run | 1, 38 | 0.56 | 0.458 | 0.002 | 0.392 |
| Dorsal premotor area | Group | 1, 38 | 6.94 | 0.012* | 0.139 | 4.138 |
| | Training run | 1, 38 | 2.80 | 0.103 | 0.008 | 0.741 |
| | Group x run | 1, 38 | 0.01 | 0.907 | < 0.001 | 0.286 |
| Primary somatosensory cortex | Group | 1, 38 | 17.54 | < 0.001* | 0.270 | 145.851 |
| | Training run | 1, 38 | 0.02 | 0.876 | < 0.001 | 0.243 |
| | Group x run | 1, 38 | 1.00 | 0.323 | 0.005 | 0.469 |

Statistical output of the group x training run ANOVAs assessing inter-block persistence in exploratory brain regions. P-values are not corrected for multiple comparisons. Significant values are marked with an asterisk, and non-significant trends are marked with a plus sign. n = 17 children, 23 adults. Corresponding results are depicted in Figure S3 below. Children exhibited larger inter-block persistence in the caudate nucleus, primary motor cortex, pre-supplementary motor area, and ventral and dorsal premotor areas as well as the primary somatosensory cortex.

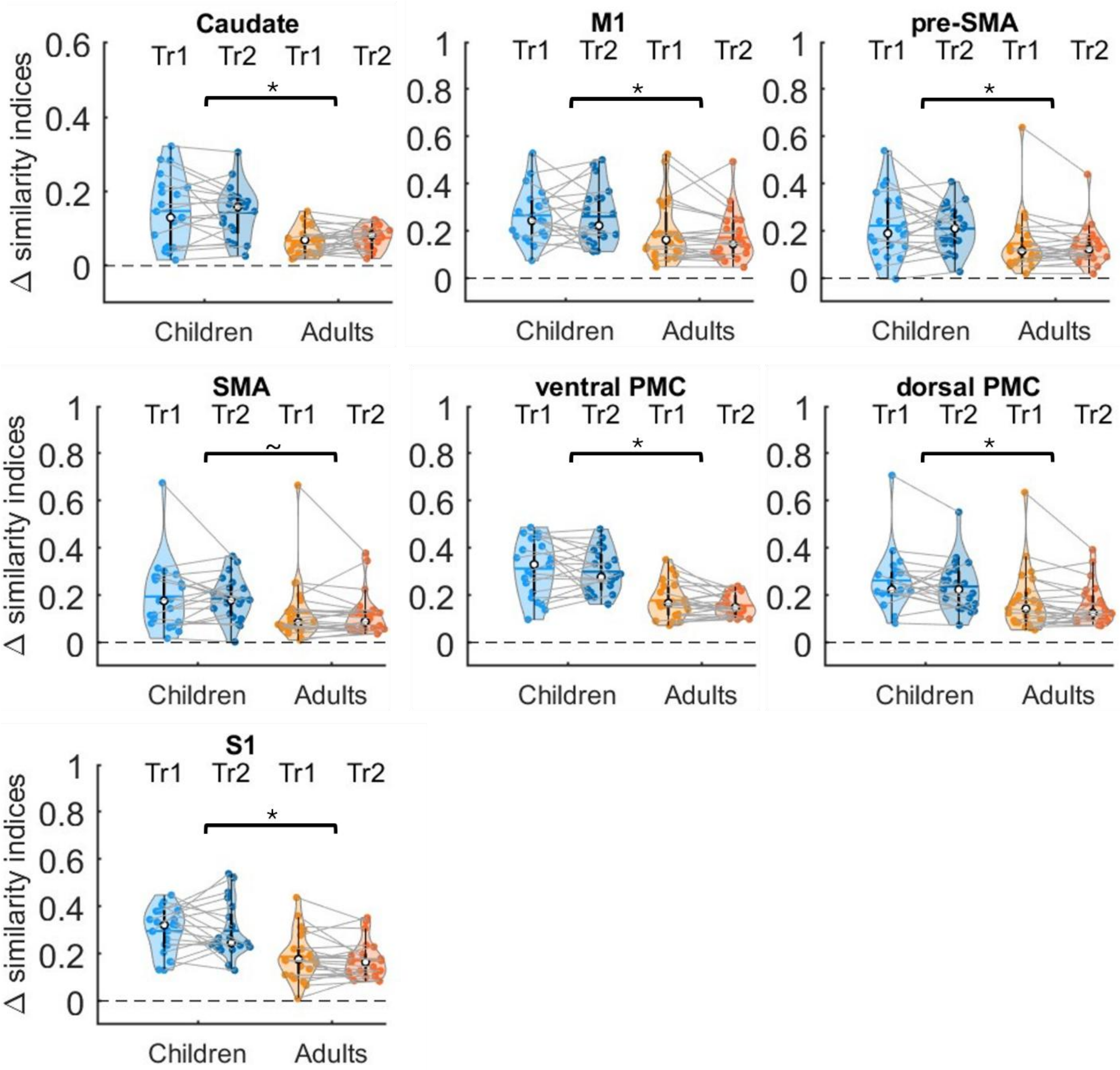

**Figure S3.** Normalized inter-block persistence for exploratory brain regions. Positive values indicate a larger SI between online task practice and inter-block rest as compared to RS1. Shaded regions represent the kernel density estimates of the data, colored circles depict individual data, open circles represent group medians, and the horizontal lines depict group means (Bechtold et al., 2021). Tr1 = training 1; n = 18 children. Tr2 = training 2; n = 19 children, unless for the supplementary motor area, as no results could be obtained for one child. n = 23 adults. * indicates $p_{uncorrected} < 0.05$ and ~ indicates $0.05 < p_{uncorrected} < 0.1$ for group effects. Statistics of group by training run ANOVAs are provided in Table S7.

**Table S8**. Statistics of linear regression analyses assessing the relationship between inter-block persistence and micro-offline performance changes in exploratory brain regions.

| Variable | | b | p |
|---|---|---|---|
| *A. Training 1* | | | |
| Caudate nucleus | Across groups | -0.257 | 0.450 |
| | Group x SI | -1.123 | 0.268 |
| Primary motor cortex | Across groups | -0.068 | 0.743 |
| | Group x SI | -0.126 | 0.778 |
| Pre-supplementary motor area | Across groups | -0.092 | 0.612 |
| | Group x SI | -0.126 | 0.739 |
| Supplementary motor area | Across groups | -0.181 | 0.360 |
| | Group x SI | -0.567 | 0.172 |
| Ventral premotor area | Across groups | 0.053 | 0.815 |
| | Group x SI | -0.561 | 0.334 |
| Dorsal premotor area | Across groups | -0.0629 | 0.744 |
| | Group x SI | -0.175 | 0.677 |
| Primary somatosensory cortex | Across groups | 0.047 | 0.843 |
| | Group x SI | 0.001 | 0.999 |
| *B. Training 2* | | | |
| Caudate nucleus | Across groups | 0.109 | 0.767 |
| | Group x SI | -0.396 | 0.698 |
| Primary motor cortex | Across groups | 0.063 | 0.731 |
| | Group x SI | 0.188 | 0.640 |
| Pre-supplementary motor area | Across groups | 0.183 | 0.406 |
| | Group x SI | 0.530 | 0.257 |
| Supplementary motor area | Across groups | 0.092 | 0.670 |
| | Group x SI | 0.367 | 0.441 |
| Ventral premotor area | Across groups | 0.055 | 0.798 |
| | Group x SI | 0.266 | 0.698 |
| Dorsal premotor area | Across groups | 0.295 | 0.137 |
| | Group x SI | 0.435 | 0.316 |
| Primary somatosensory cortex | Across groups | -0.023 | 0.908 |
| | Group x SI | 0.200 | 0.685 |

Training 1: n = 23 adults, 18 children. Training 2: n = 23 adults, 19 children, unless for the supplementary motor area, as no results could be obtained for one child. No significant relationships were observed. P-values are not corrected for multiple comparisons.

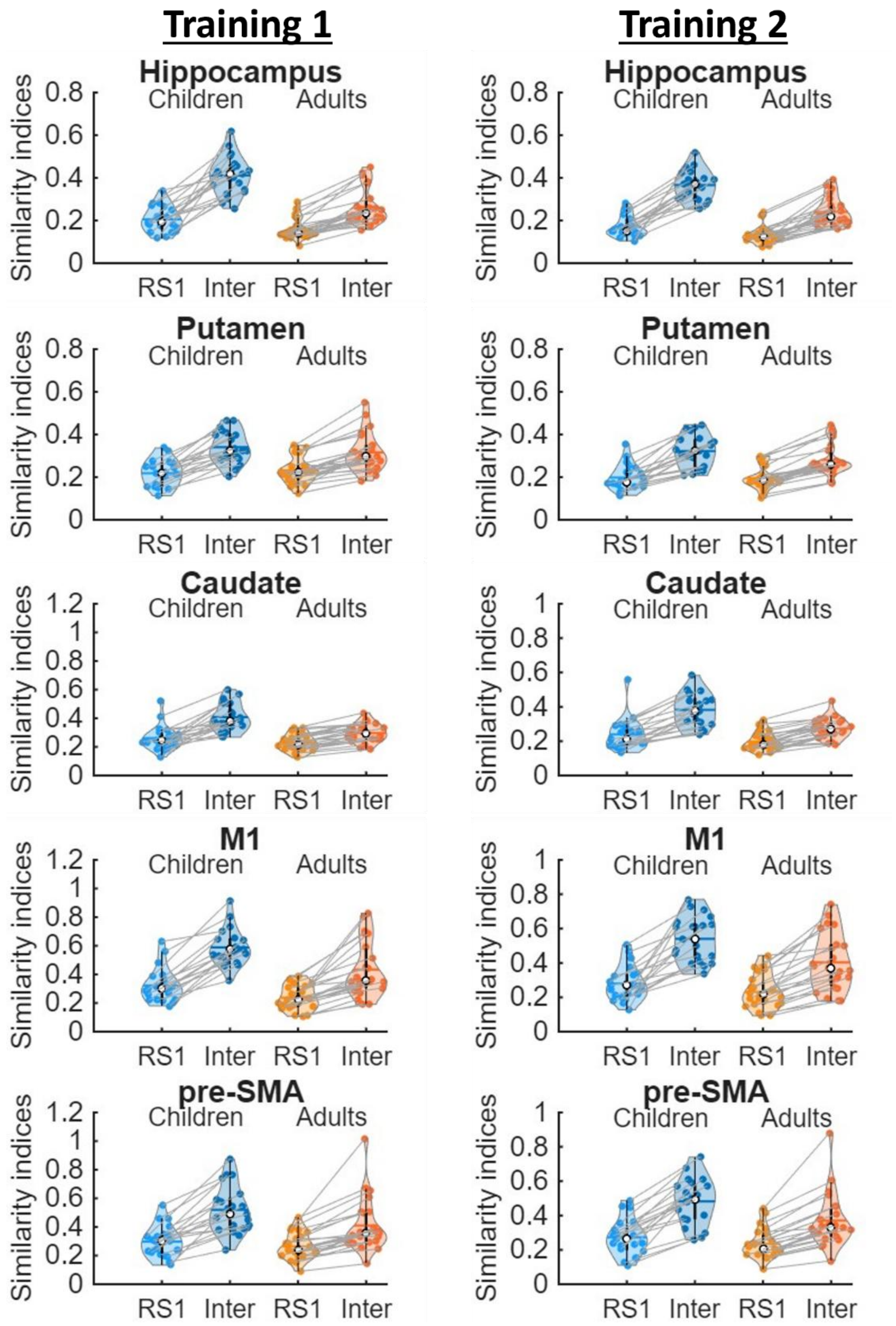

**Figure S4.** Similarity of task activity patterns during training with patterns of activity during pre-learning (i.e., RS1) and inter-block rest. Similarity indices were computed using r-to-z transformed correlations. Shaded regions represent the kernel density estimates of the data, colored circles depict individual data, open circles represent group medians, and the horizontal lines depict group means (Bechtold et al., 2021). Training 1 = 18 children, Training 2 = 19 children, unless for the supplementary motor area, as no results could be obtained for one child. n = 23 adults.

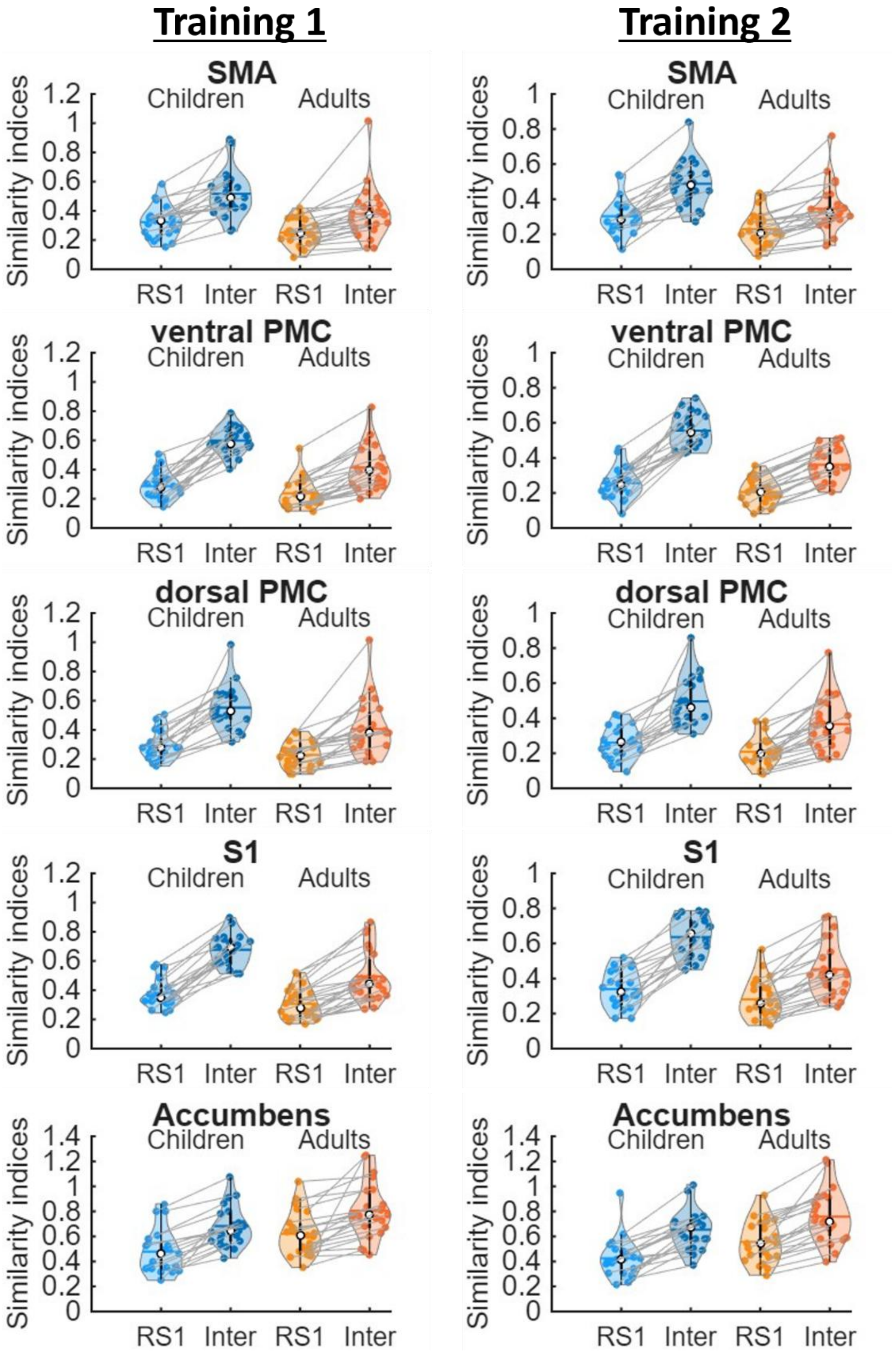

**Figure S4 continued.** Similarity of task activity patterns during training with patterns of activity during pre-learning (i.e., RS1) and inter-block rest. Similarity indices were computed using r-to-z transformed correlations. Shaded regions represent the kernel density estimates of the data, colored circles depict individual data, open circles represent group medians, and the horizontal lines depict group means (Bechtold et al., 2021). Training 1 = 18 children, Training 2 = 19 children, unless for the supplementary motor area, as no results could be obtained for one child. n = 23 adults.

##### 4.2. Persistence into post-learning rest

**Table S9.** Results of regressions that tested the relationship between age and post-learning persistence within the sample of children.

| | **Post-learning Persistence** | |
|---|---|---|
| **Region of interest** | **Training 1** | **Training 2** |
| Hippocampus | $R^2$ = 0.017; p = 0.405 | $R^2$ = 0.022; p = 0.754 |
| Putamen | $R^2$ = 0.043; p = 0.753 | $R^2$ = 0.013; p = 0.827 |
| Nucleus accumbens | $R^2$ = 0.019; p = 0.494 | $R^2$ = 0.019; p = 0.713 |
| Caudate nucleus | $R^2$ = 0.073; p = 0.225 | $R^2$ = 0.003; p = 0.926 |
| Primary motor cortex | $R^2$ = 0.027; p = 0.478 | $R^2$ = 0.003; p = 0.812 |
| Pre-supplementary motor area | $R^2$ = 0.061; p = 0.292 | $R^2$ = 0.061; p = 0.294 |
| Supplementary motor area | $R^2$ = 0.110; p = 0.141 | $R^2$ = 0.079; p = 0.217 |
| Ventral premotor area | $R^2$ = 0.071; p = 0.244 | $R^2$ = 0.018; p = 0.561 |
| Dorsal premotor area | $R^2$ = 0.076; p = 0.226 | $R^2$ = 0.1151 p = 0.133 |
| Primary somatosensory cortex | $R^2$ = 0.065; p = 0.267 | $R^2$ = 0.005; p = 0.771 |

$R^2$ = R-squared. No significant relationships were observed. P-values are uncorrected for multiple comparisons. Results are visualized in S5-S6 below.

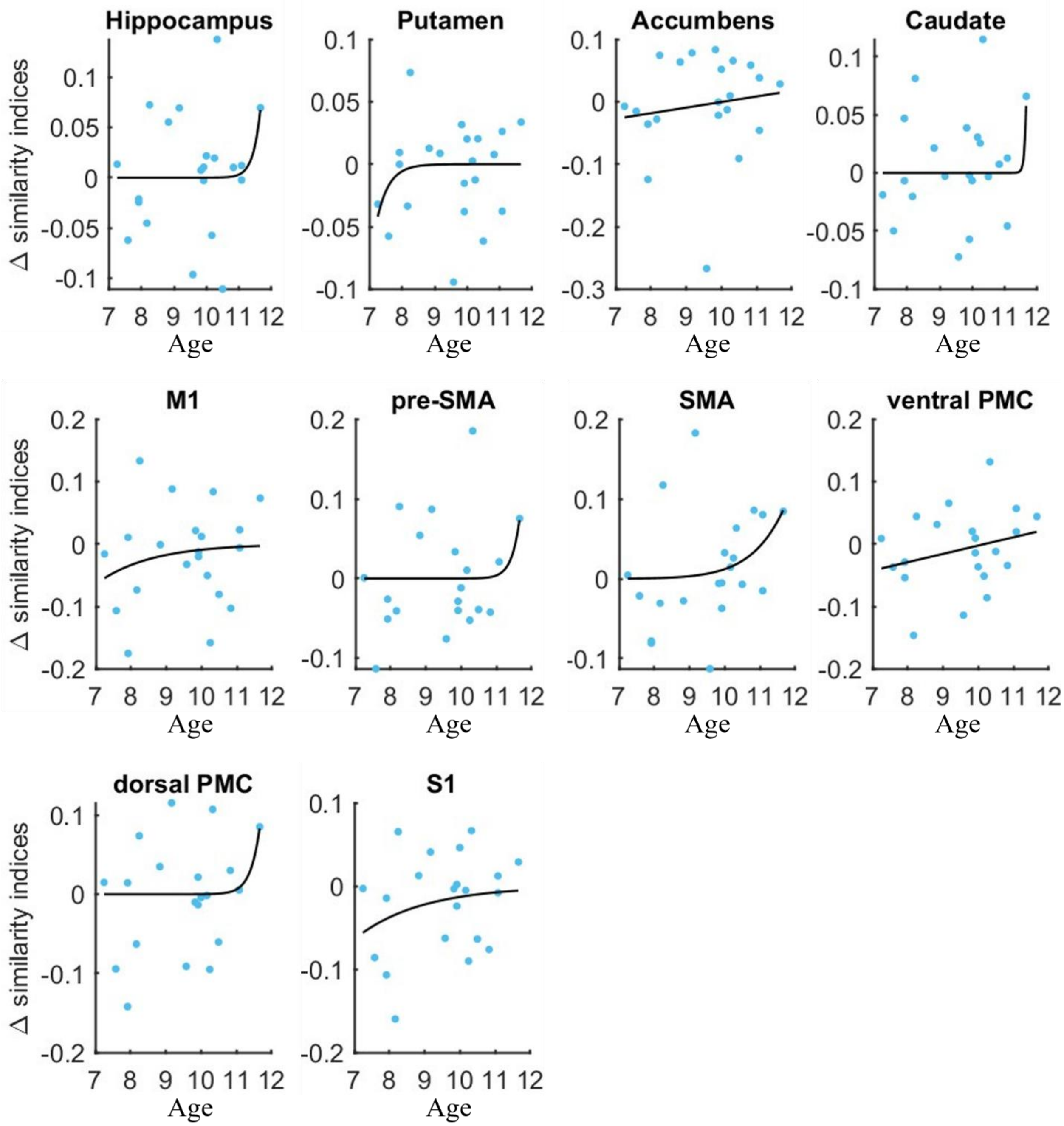

**Figure S5.** Post-learning persistence of task activity during training 1 plotted as a function of age within children for primary and exploratory regions. Potential fit options were single exponential, double exponential, linear, quadratic and power functions. M1 = primary motor cortex, pre-SMA = pre-supplementary motor area, SMA = supplementary motor area, PMC = premotor cortex, S1 = primary somatosensory cortex. Statistics are provided in Supplementary Table S9 above. None of the relationships were significant at $p < 0.05$.

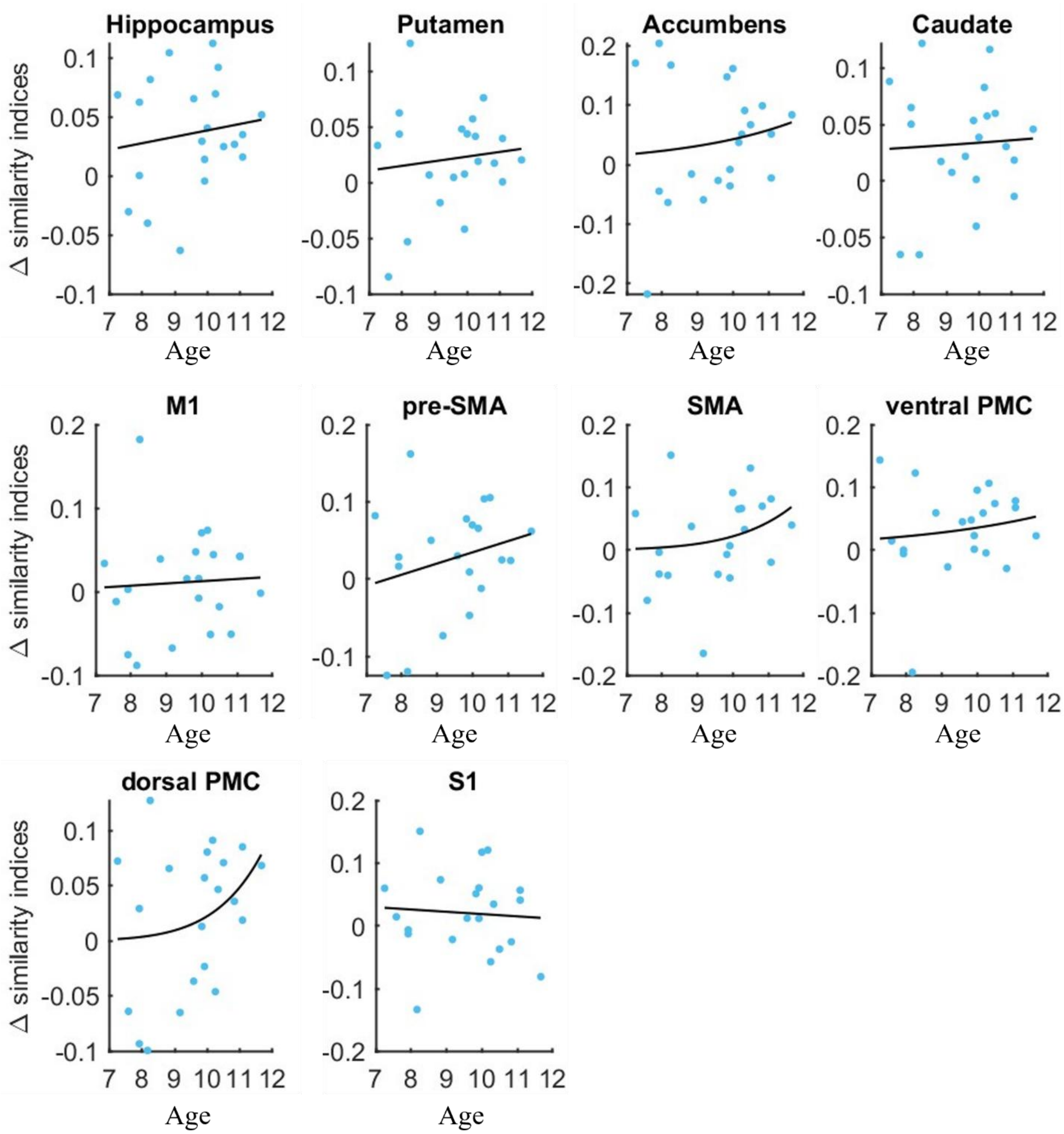

**Figure S6.** Post-learning persistence of task activity during training 2 plotted as a function of age within children for primary and exploratory regions. Potential fit options were single exponential, double exponential, linear, quadratic and power functions. M1 = primary motor cortex, pre-SMA = pre-supplementary motor area, SMA = supplementary motor area, PMC = premotor cortex, S1 = primary somatosensory cortex. Statistics are provided in Supplementary Table S9. None of the relationships were significant at $p < 0.05$.

**Table S10.** Results of group x run ANOVAs on post-learning persistence in exploratory regions.

| Region | Effect | df | F | p | $\eta^2$ | $BF_{10}$ |
|---|---|---|---|---|---|---|
| Caudate nucleus | Group | 1, 41 | 2.55 | 0.118 | 0.052 | 0.842 |
| | Training run | 1, 41 | 38.72 | < 0.001* | 0.097 | 5.37*e$^4$ |
| | Group x run | 1, 41 | 1.07 | 0.307 | 0.003 | 0.469 |
| Primary motor cortex | Group | 1, 41 | 0.01 | 0.918 | < 0.001 | 0.351 |
| | Training run | 1, 41 | 4.89 | 0.033* | 0.025 | 1.311 |
| | Group x run | 1, 41 | 2.88 | 0.097$^+$ | 0.015 | 0.881 |
| Pre-supplementary motor area | Group | 1, 40 | 1.16 | 0.288 | 0.017 | 0.407 |
| | Training run | 1, 40 | 2.70 | 0.108 | 0.026 | 0.764 |
| | Group x run | 1, 40 | 0.86 | 0.359 | 0.008 | 0.458 |
| Supplementary motor area | Group | 1, 41 | 1.17 | 0.285 | 0.017 | 0.402 |
| | Training run | 1, 41 | 0.72 | 0.400 | 0.007 | 0.347 |
| | Group x run | 1, 41 | 0.07 | 0.797 | < 0.001 | 0.318 |
| Ventral premotor area | Group | 1, 41 | 2.18 | 0.147 | 0.038 | 0.508 |
| | Training run | 1, 41 | 19.77 | < 0.001* | 0.108 | 282.546 |
| | Group x run | 1, 41 | 3.54 | 0.067$^+$ | 0.021 | 1.644 |
| Dorsal premotor area | Group | 1, 41 | 0.61 | 0.437 | 0.011 | 0.361 |
| | Training run | 1, 41 | 3.87 | 0.056$^+$ | 0.027 | 1.202 |
| | Group x run | 1, 41 | 1.38 | 0.247 | 0.010 | 0.614 |
| Primary sensory cortex | Group | 1, 41 | 0.02 | 0.885 | < 0.001 | 0.316 |
| | Training run | 1, 41 | 12.32 | 0.001* | 0.076 | 31.107 |
| | Group x run | 1, 41 | 2.80 | 0.102 | 0.018 | 1.195 |

Statistical output of the group x training run ANOVAs assessing post-learning persistence in exploratory brain regions. Corresponding results are depicted in Figure S7 below. P-values are not corrected for multiple comparisons. Significant values are marked with an asterisk, and non-significant trends are marked with a plus sign. n = 20 children, 23 adults.

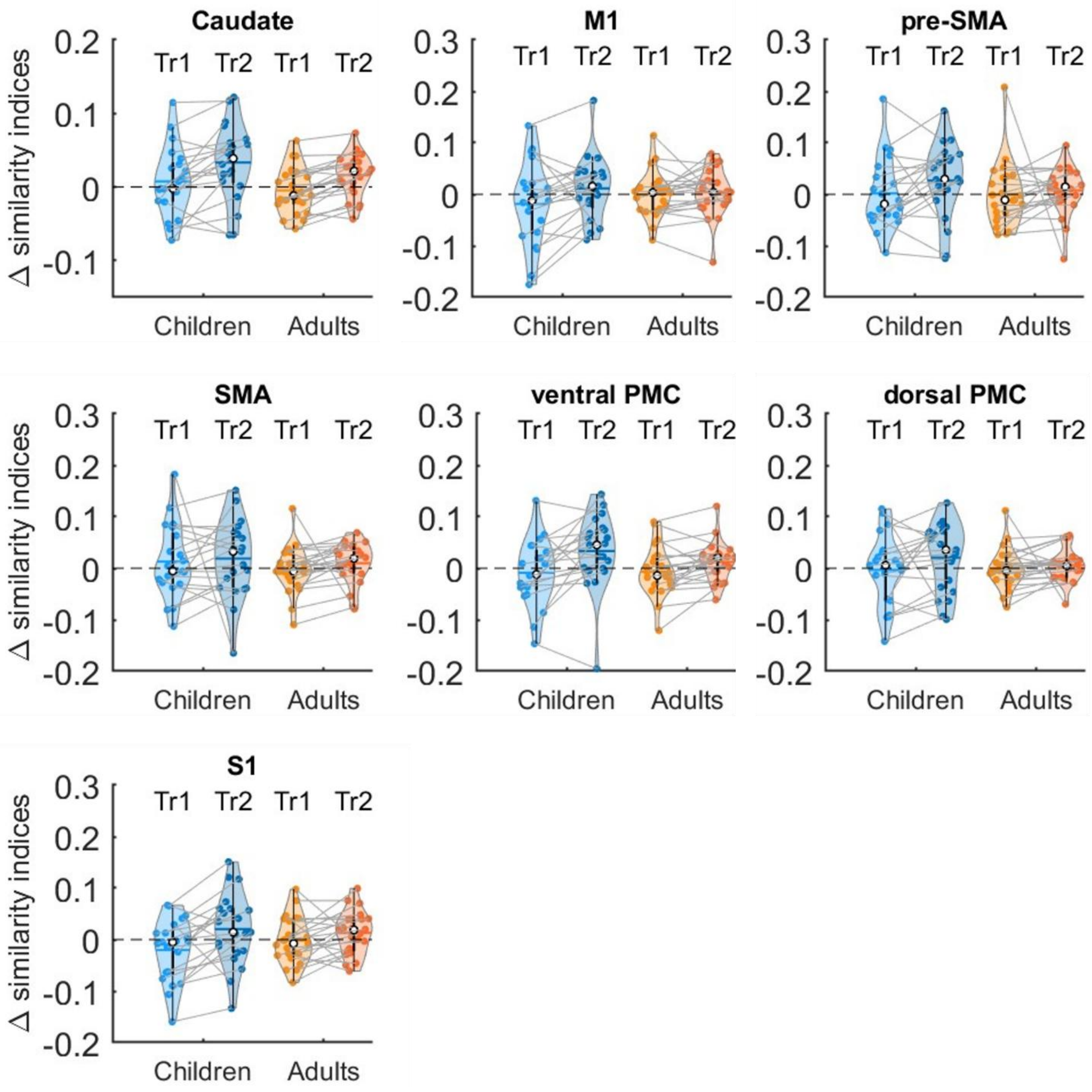

**Figure S7**. Post-learning persistence for exploratory brain regions. Positive values indicate a larger SI between online task practice and RS2 as compared RS1. Shaded regions represent the kernel density estimates of the data, colored circles depict individual data, open circles represent group medians, and the horizontal lines depict group means (Bechtold et al., 2021). n = 20 children and 23 adults. Results of group by training run ANOVAs are depicted in Table S10.

**Table S11.** Statistics of linear regression analyses assessing the relationships between post-learning persistence and macro-offline performance changes in exploratory brain regions.

| Variable | | b | p |
|---|---|---|---|
| *A. Training 1* | | | |
| Caudate nucleus | Across groups | 0.258 | 0.528 |
| | Group x SI | -0.760 | 0.389 |
| Primary motor cortex | Across groups | 0.004 | 0.989 |
| | Group x SI | 0.594 | 0.321 |
| Pre-supplementary motor area | Across groups | -0.040 | 0.884 |
| | Group x SI | 0.189 | 0756 |
| Supplementary motor area | Across groups | 0.082 | 0.744 |
| | Group x SI | 0.290 | 0.570 |
| Ventral premotor area | Across groups | 0.044 | 0.880 |
| | Group x SI | 0.645 | 0.280 |
| Dorsal premotor area | Across groups | -0.046 | 0.875 |
| | Group x SI | 1.044 | 0.115 |
| Primary sensory cortex | Across groups | -0.201 | 0.507 |
| | Group x SI | 0.988 | 0.116 |
| *B. Training 2* | | | |
| Caudate nucleus | Across groups | -0.024 | 0.953 |
| | Group x SI | -0.147 | 0.876 |
| Primary motor cortex | Across groups | -0.247 | 0.425 |
| | Group x SI | 0.338 | 0.596 |
| Pre-supplementary motor area | Across groups | -0.165 | 0.551 |
| | Group x SI | 0.094 | 0.886 |
| Supplementary motor area | Across groups | -0.026 | 0.929 |
| | Group x SI | 0.024 | 0.970 |
| Ventral premotor area | Across groups | -0.138 | 0.692 |
| | Group x SI | 1.472 | $0.050^{+}$ |
| Dorsal premotor area | Across groups | -0.104 | 0.753 |
| | Group x SI | 0.736 | 0.349 |
| Primary sensory cortex | Across groups | -0.263 | 0.402 |
| | Group x SI | 0.944 | 0.153 |

Training 1: n = 21 children. Training 2: n = 20 children, unless for the supplementary motor area, as no results could be obtained for one child. n = 23 adults. SI = similarity index. P-values are uncorrected for multiple comparisons. $^{+}$ corresponds to a p-value less than 0.1.

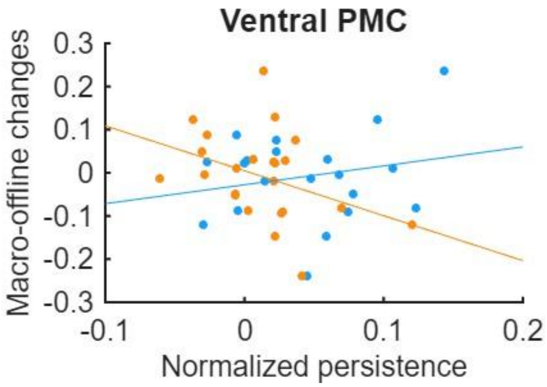

**Figure S8**. Relationships between macro-offline performance changes and post-learning persistence for the ventral premotor cortex (PMC) for children (blue) and adults (orange). Positive persistence values indicate a larger SI between online task practice and RS2 as compared RS1. Positive macro-offline changes indicate improvements from the post-learning test to 5-hour retest. Colored circles depict individual datapoints. n = 20 children and 23 adults. Results of regression analyses are depicted in Table S11.

**Table S12.** Results of group x RS ANOVAs comparing persistence into the post-consolidation RS3 vs. persistence into RS2.

| Region | Effect | df | F | p | $\eta^2$ | $BF_{10}$ |
|---|---|---|---|---|---|---|
| Hippocampus | Group | 1, 42 | 20.78 | < 0.001* | 0.251 | 45.545 |
| | RS run | 2, 83.85 | 192.16 | < 0.001* | 0.595 | $3.439*e^{28}$ |
| | Group x RS | 2, 83.85 | 4.39 | 0.015* | 0.033 | 3.035 |
| Putamen | Group | 1, 42 | 0.40 | 0.530 | 0.007 | 0.292 |
| | RS run | 1.87, 78.54 | 145.69 | < 0.001* | 0.421 | $1.684*e^{25}$ |
| | Group x RS | 1.87, 78.54 | 1.13 | 0.326 | 0.006 | 0.308 |
| Accumbens | Group | 1, 42 | 4.81 | 0.034* | 0.055 | 0.477 |
| | RS run | 1.25, 52.39 | 169.96 | < 0.001* | 0.666 | $1.987*e^{29}$ |
| | Group x RS | 1.25, 52.39 | 10.15 | 0.001* | 0.107 | 289.598 |
| Caudate nucleus | Group | 1, 42 | 7.05 | 0.011* | 0.119 | 3.936 |
| | RS run | 1.76, 73.93 | 138.29 | < 0.001* | 0.390 | $1.516*e^{24}$ |
| | Group x RS | 1.76, 73.93 | 0.94 | 0.385 | 0.004 | 0.266 |
| Primary motor cortex | Group | 1, 42 | 3.09 | $0.086^{+}$ | 0.057 | 0.997 |
| | RS run | 1.82, 76.48 | 89.07 | < 0.001* | 0.281 | $1.048*e^{18}$ |
| | Group x RS | 1.82, 76.48 | 1.86 | 0.165 | 0.008 | 0.458 |
| Pre-supplementary motor area | Group | 1, 42 | 8.28 | 0.006* | 0.130 | 6.686 |
| | RS run | 1.75, 73.56 | 70.47 | < 0.001* | 0.287 | $1.855*e^{15}$ |
| | Group x RS | 1.75, 73.56 | 2.02 | 0.145 | 0.011 | 0.548 |
| Supplementary motor area | Group | 1, 41 | 4.12 | 0.049* | 0.069 | 0.904 |
| | RS run | 1.81, 74.04 | 76.82 | < 0.001* | 0.331 | $1.145*e^{16}$ |
| | Group x RS | 1.81, 74.04 | 1.74 | 0.186 | 0.011 | 0.390 |
| Ventral premotor area | Group | 1, 42 | 5.65 | 0.022* | 0.098 | 2.363 |
| | RS run | 1.97, 82.87 | 128.93 | < 0.001* | 0.368 | $2.570*e^{22}$ |
| | Group x RS | 1.97, 82.87 | 3.14 | 0.049* | 0.014 | 1.220 |
| Dorsal premotor area | Group | 1, 42 | 6.02 | 0.018* | 0.103 | 3.099 |
| | RS run | 1.82, 76.42 | 80.06 | < 0.001* | 0.273 | $1.061*e^{17}$ |
| | Group x RS | 1.82, 76.42 | 1.01 | 0.364 | 0.005 | 0.241 |
| Primary sensory cortex | Group | 1, 42 | 3.06 | $0.088^{+}$ | 0.058 | 0.992 |
| | RS run | 1.68, 70.68 | 126.93 | < 0.001* | 0.310 | $2.381*e^{22}$ |
| | Group x RS | 1.68, 70.68 | 2.38 | 0.109 | 0.008 | 0.703 |

Statistical output of the group x RS run ANOVAs assessing post-task persistence in both primary and exploratory brain regions. P-values are not corrected for multiple comparisons. Significant values are marked with an asterisk, non-significant trends are marked with a plus sign. n = 20 children, except for the supplementary motor area, as no results could be obtained for one child. n = 23 adults.

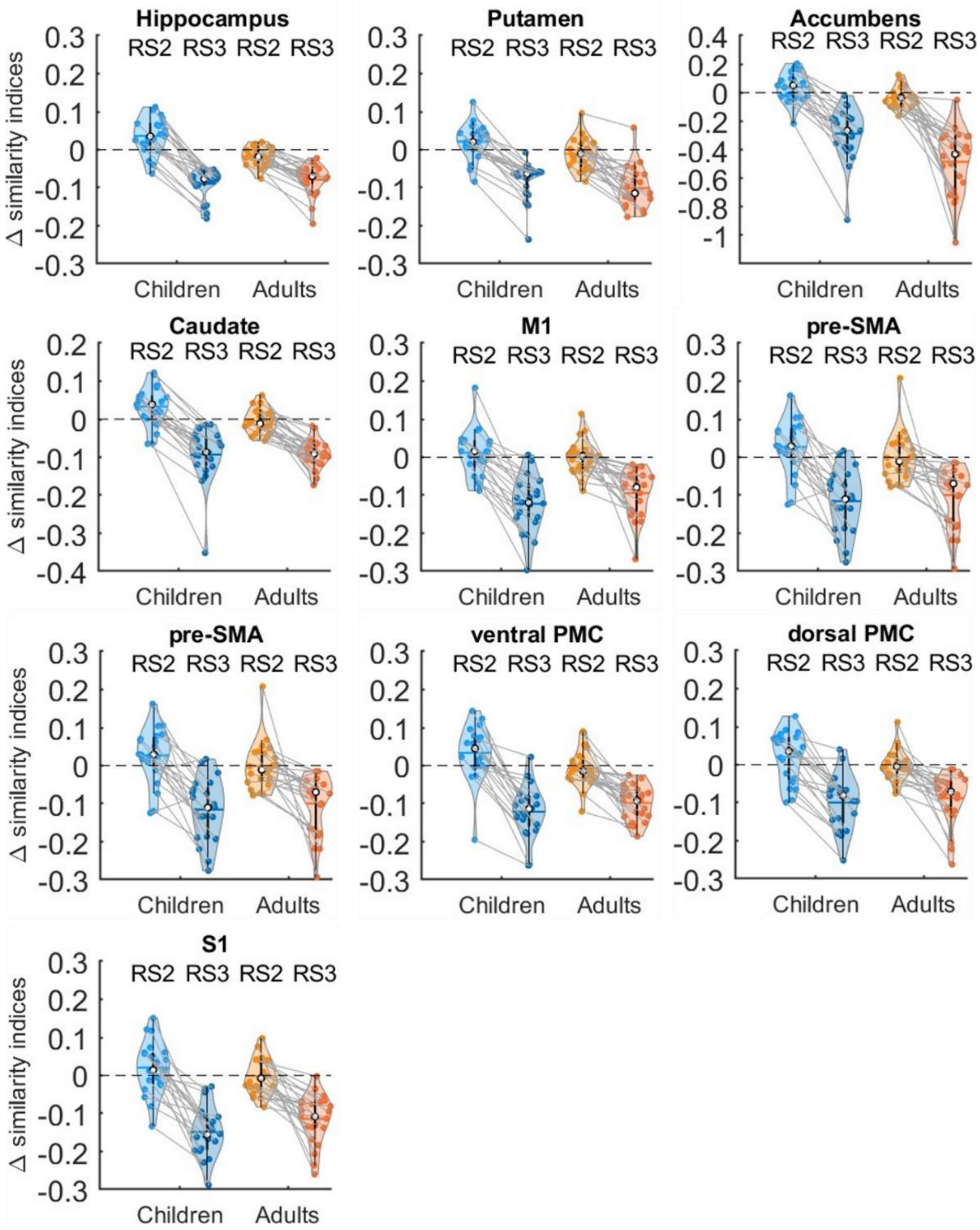

**Figure S9.** Persistence of task activity during training 2 into post-learning (i.e., RS2) and post-consolidation rest (i.e., RS3). Similarity indices (SI) between online task practice of training 2 and the pre-, post-learning and post-consolidation RS runs were computed using r-to-z transformed correlations. Normalized SIs were obtained by subtracting the SIs between online task practice and RS1 from the SIs between online task practice and RS2 and RS3. Positive values indicate a larger SI between online task practice and RS2 or RS3 as compared RS1. Shaded regions represent the kernel density estimates of the data, colored circles depict individual data, open circles represent group medians, and the horizontal lines depict group means (Bechtold et al., 2021). n = 20 children and 23 adults. Statistics of group by RS run ANOVAs are depicted in Table S12.

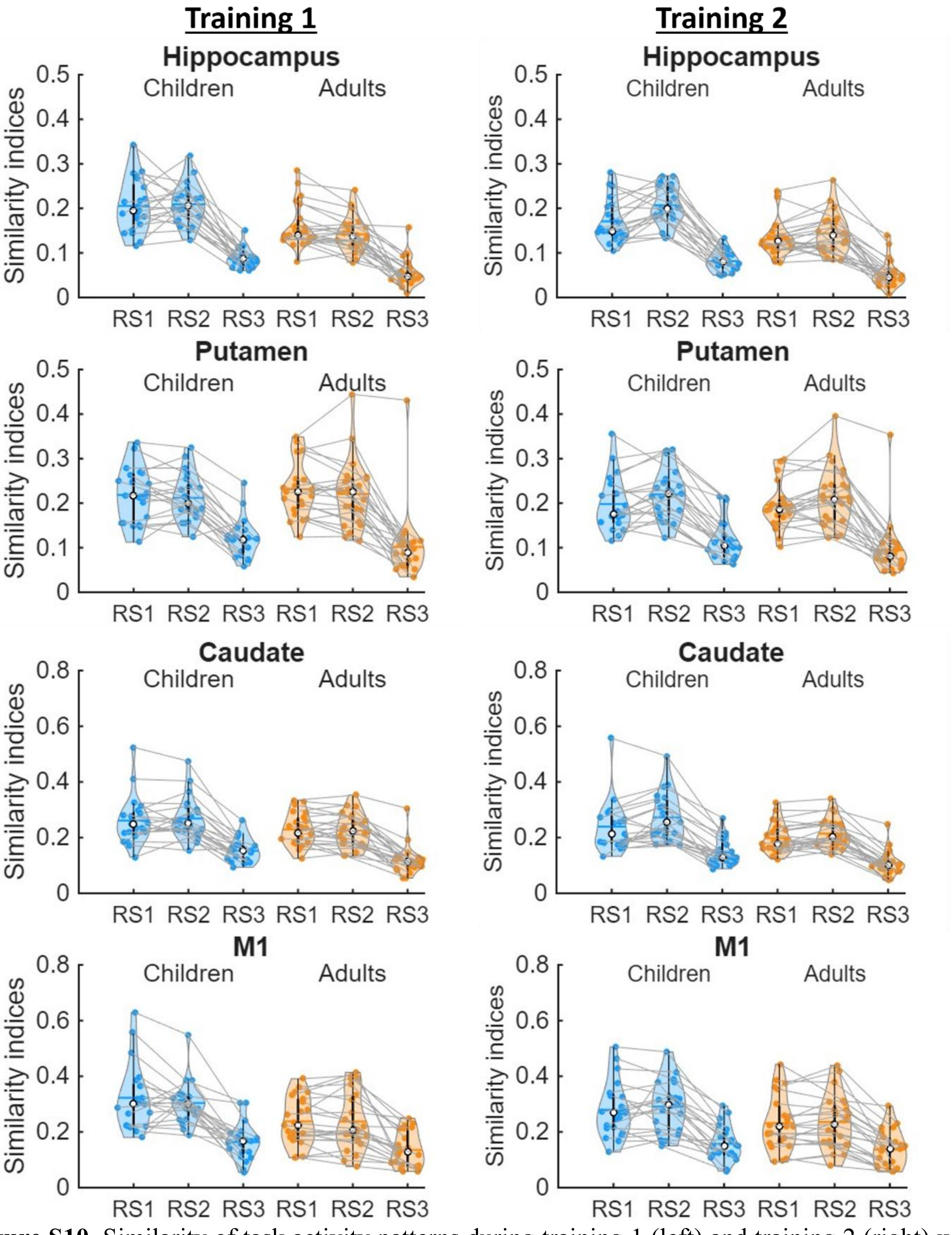

**Figure S10.** Similarity of task activity patterns during training 1 (left) and training 2 (right) with patterns of activity during pre-learning (i.e., RS1), post-learning (i.e., RS2) and post-consolidation rest (i.e., RS3). Similarity indices were computed using r-to-z transformed correlations. Shaded regions represent the kernel density estimates of the data, colored circles depict individual data, open circles represent group medians, and the horizontal lines depict group means (Bechtold et al., 2021). n = 20 children and 23 adults.

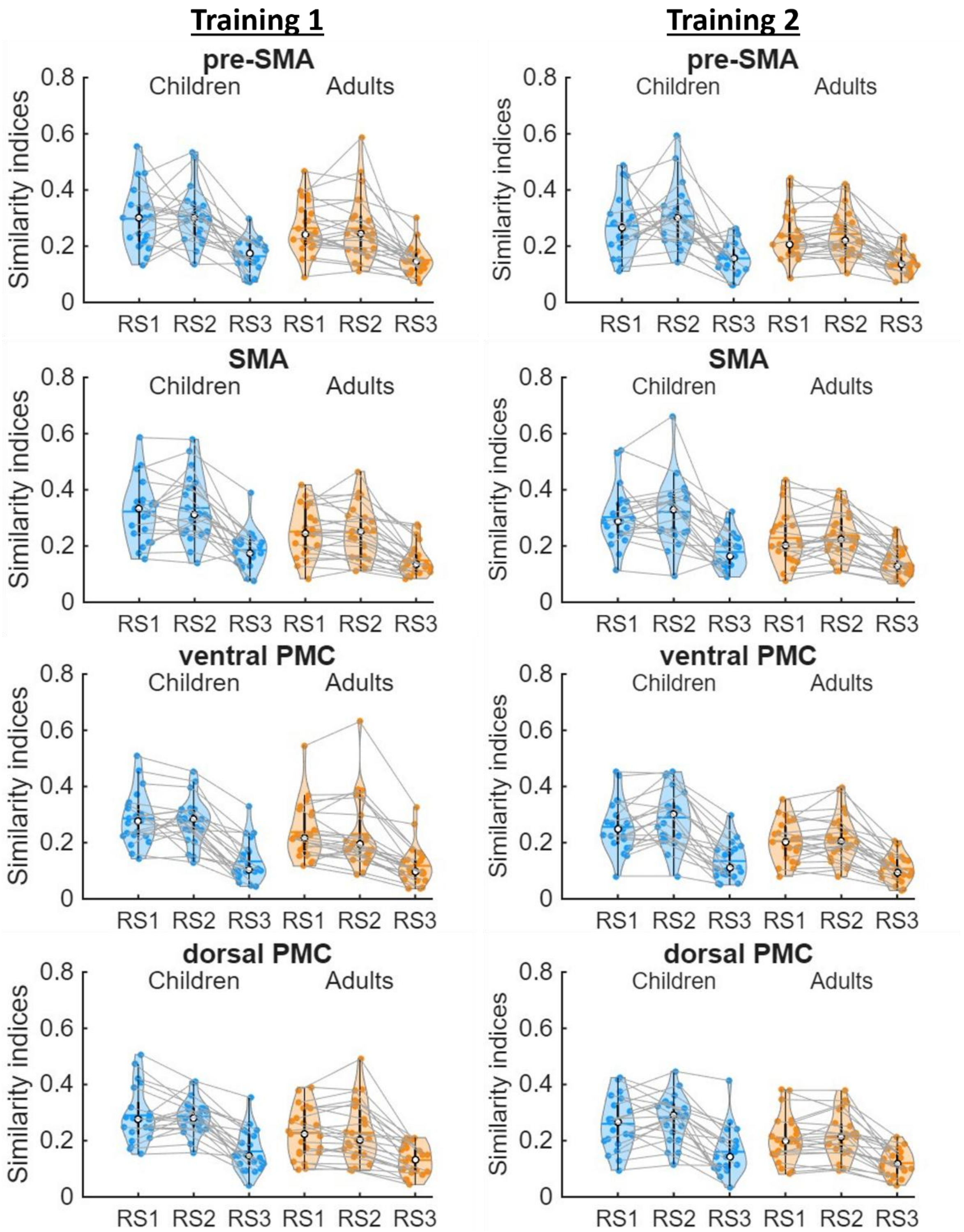

**Figure S10 continued.** Similarity of task activity patterns during training 1 (left) and training 2 (right) with patterns of activity during pre-learning (i.e., RS1), post-learning (i.e., RS2) and post-consolidation rest (i.e., RS3). Similarity indices were computed using r-to-z transformed correlations. Shaded regions represent the kernel density estimates of the data, colored circles depict individual data, open circles represent group medians, and the horizontal lines depict group means (Bechtold et al., 2021). n = 20 children and 23 adults.

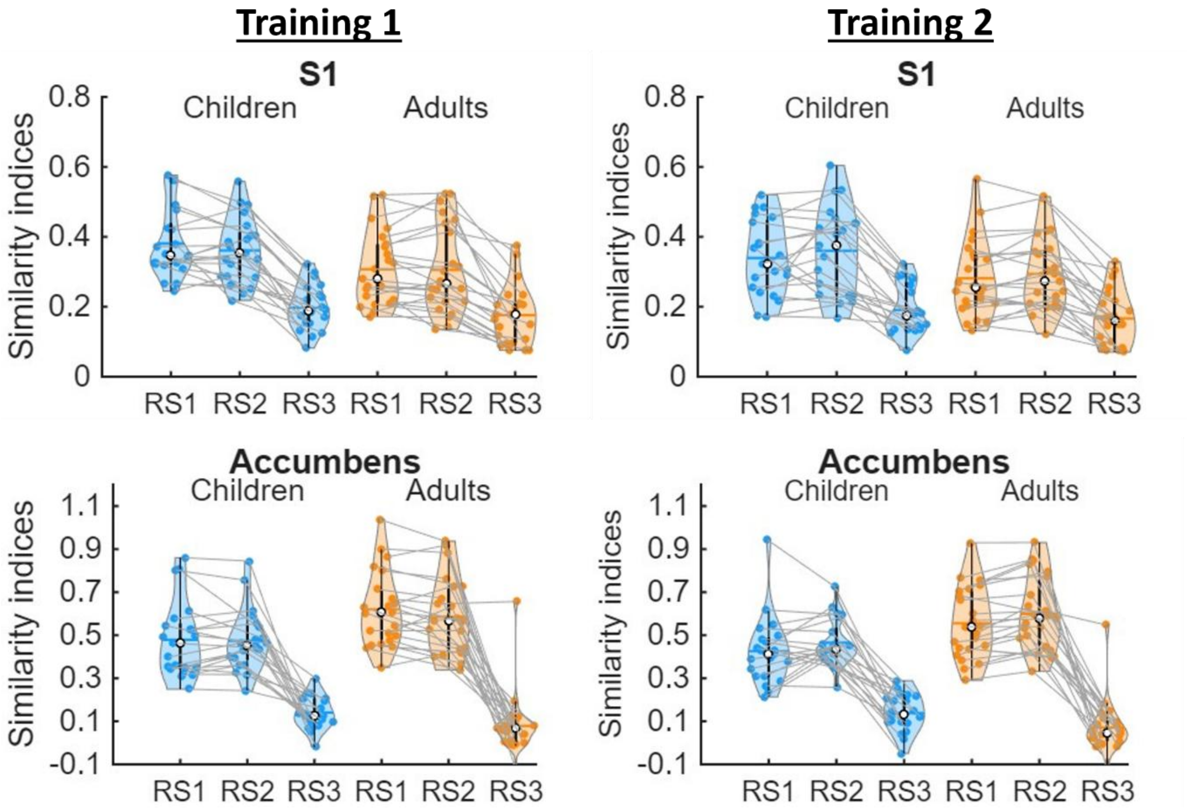

**Figure S10 continued.** Similarity of task activity patterns during training with patterns of activity during pre-learning (i.e., RS1), post-learning (i.e., RS2) and post-consolidation rest (i.e., RS3). Similarity indices were computed using r-to-z transformed correlations. Shaded regions represent the kernel density estimates of the data, colored circles depict individual data, open circles represent group medians, and the horizontal lines depict group means (Bechtold et al., 2021). n = 20 children and 23 adults.
